# Full-length annotation with multi-strategy RNA-seq uncovers transcriptional regulation of lncRNAs in diploid cotton *G. arboreum*^1^

**DOI:** 10.1101/2020.07.21.214502

**Authors:** Xiaomin Zheng, Yanjun Chen, Yifan Zhou, Danyang Li, Keke Shi, Xiao Hu, Hanzhe Ye, Yu Zhou, Kun Wang

## Abstract

Long noncoding RNAs (lncRNAs) are crucial factors during plant development and environmental responses. High-throughput and accurate identification of lncRNAs is still lacking in plants. To build an accurate atlas of lncRNA in cotton, we combined Isoform-sequencing (Iso-seq), strand-specific RNA-seq (ssRNA-seq), cap analysis gene expression (CAGE-seq) with PolyA-seq and compiled a pipeline named plant full-length lncRNA (PULL) to integrate multi-omics data. A total of 9240 lncRNAs from 21 tissue samples of the diploid cotton *Gossypium arboreum* were identified. We revealed that alternative usage of transcription start site (TSS) and transcription end site (TES) of lncRNAs occurs pervasively during plant growth and responses to stress. We identified the lncRNAs which co-expressed or be linked to the protein coding genes (PCGs) or GWAS studied SNPs associated with ovule and fiber development. We also mapped the genome-wide binding sites of two lncRNAs with chromatin isolation by RNA purification sequencing (ChIRP-seq) and validated the *trans* transcriptional regulation of *lnc-Ga13g0352* via virus induced gene suppression (VIGS) assay. These findings provide valuable research resources for plant community and broaden our understandings of biogenesis and regulation function of plant lncRNAs.

**One sentence summary:** The full-length annotation and transcriptional regulation of long noncoding RNAs in cotton.

## INTRODUCTION

Long noncoding RNAs (LncRNAs) have been proved to play an essential role in gene transcriptional regulation. In plants, *AUXIN REGULATED PROMOTER LOOP RNA* (*APOLO*) acts as a scaffold to control chromatin looping and DNA methylation (Ariel et al., 2014); The antisense lncRNA *Cold Induced Long Antisense Intragenic RNA* (*COOLAIR)* mediates chromatin switching at *FLOWERING LOCUS C* (*FLC*) during vernalization (Csorba et al., 2014); *INDUCED BY PHOSPHATE STARVATION1/2* (*IPS1/2*) sponges *miR399* to regulate its target, *PHOSPHATE2* (*PHO2*), and maintain correct *Pi* homeostasis (Franco-Zorrilla et al., 2007); *Long-day-specific Male-fertility Associated RNA* (*LDMAR*) is required for normal pollen development by causing promoter methylation under long-day conditions (Ding et al., 2012); *Alternative splicing competitor long noncoding RNA* (*ASCO-lncRNA*) can hijack nuclear alternative splicing (AS) regulators to modulate AS patterns during development (Bardou et al., 2014); Over-expression of *LRK Antisense Intergenic RNA* (*LAIR*) might regulate the expression of neighboring gene cluster to increase grain yield in rice (Wang et al., 2018b).

However, the gene structures, biogenesis, expression and molecular functions of plant lncRNAs are still largely under exploration (St.Laurent et al., 2015). Currently, the saturated annotation resources for lncRNAs are limited to few model plants, such as *Arabidopsis thaliana* (Liu et al., 2012), maize (Li et al., 2014b) and rice (Zhang et al., 2014). Although more than 12,000 lncRNAs have been deposited in the GreeNC database (Paytuví Gallart et al., 2016), most lncRNAs in the database were identified only by using Illumina next generation sequencing (NGS) RNA-seq, in which the annotations are obtained from short reads assembly, thus producing incomplete and inaccurate annotations on lncRNA’s boundaries: junction sites and 5’/3’-ends (Boley et al., 2014)(Uszczynska-Ratajczak et al., 2018).

With the improvement in high-throughput sequencing and associated computational methods, revolutionary methods for lncRNAs identification have been constructed. The Cap analysis gene expression (CAGE-seq) and PolyA-seq capture sequencing short reads that originate from the 5’ cap and 3’ polyA tail, offering precise information of transcription start site (TSSs) and transcription end site (TESs) for RNA transcripts. PacBio long-read sequencing directly collects full-length transcripts to synchronously produce splicing, TSS and TES information. These methods have already been combined with RNA-seq for RNA annotation in humans and other animals (Boley et al., 2014) (Hon et al., 2017).

Cotton is an important cash crop worldwide. In addition to producing animal feed and cooking oil, cotton seeds are known to be the source of pure cellulose fiber. The fiber cell development of cotton is an excellent model for studying cell elongation and cell wall biosynthesis (Wang et al., 2016). To date, by using NGS RNA-seq, two studies of lncRNA identification have been performed in two tetraploid cotton species, *G. barbadense* and *G. hirsutum* (Wang et al., 2015a) (Zhao et al., 2018a).

In previous studies, we have sequenced and assembled the genome and performed transcriptomic annotation for the cultivated diploid cotton *G. arboreum* (Li et al., 2014a)(Du et al., 2018)(Wang et al., 2019a). In order to obtain systematic and accurate information of lncRNAs in *G. arboreum*, we combined multiple RNA-seq methods, including Isoform-sequencing (Iso-seq), strand-specific RNA sequencing (ssRNA-seq), CAGE-seq, and PolyA-seq to investigate the lncRNA expression in cotton (*G. arboreum*). We compiled the PULL pipeline to integrate these data and finally identified 9240 lncRNAs across 21 tissue samples. This study represents a fundamental resource to enrich the knowledge base on lncRNAs, allowing the cotton community to utilize these results in future functional studies.

## RESULTS

### Cotton lncRNA transcriptome with accurate TSSs and TESs

To obtain comprehensive annotation for lncRNAs in *G. arboreum*, we analyzed four types of sequencing data across 16 tissues and 5 stress treated samples obtained from Iso-seq, ssRNA-seq (ribosome depletion-based sequence library), CAGE-seq, and PolyA-seq (**Fig. 1A**). We compiled a pipeline, named plant full-length lncRNA (PULL) to systematically integrate multi-omics data and aimed to identify accurate gene structures, including TSSs, TESs and junction sites for lncRNAs (**Fig. 1B**; Supplemental Fig. S1). Transcriptome assembled by ssRNA-seq and Iso-seq was corrected by CAGE-seq/PolyA-seq to obtain accurate TSS/TES. The whole set of isoforms was then filtered to obtain the lncRNAs through removing protein coding genes (PCGs), known ncRNAs in Rfam (tRNAs, rRNAs, snoRNAs, and snRNAs) (Kalvari et al., 2018), lncRNAs with high score of coding potential, and possible false-positive long noncoding natural antisense transcripts (lnc-NATs) (Supplemental Fig. S1; details in methods).

**Figure 1.**
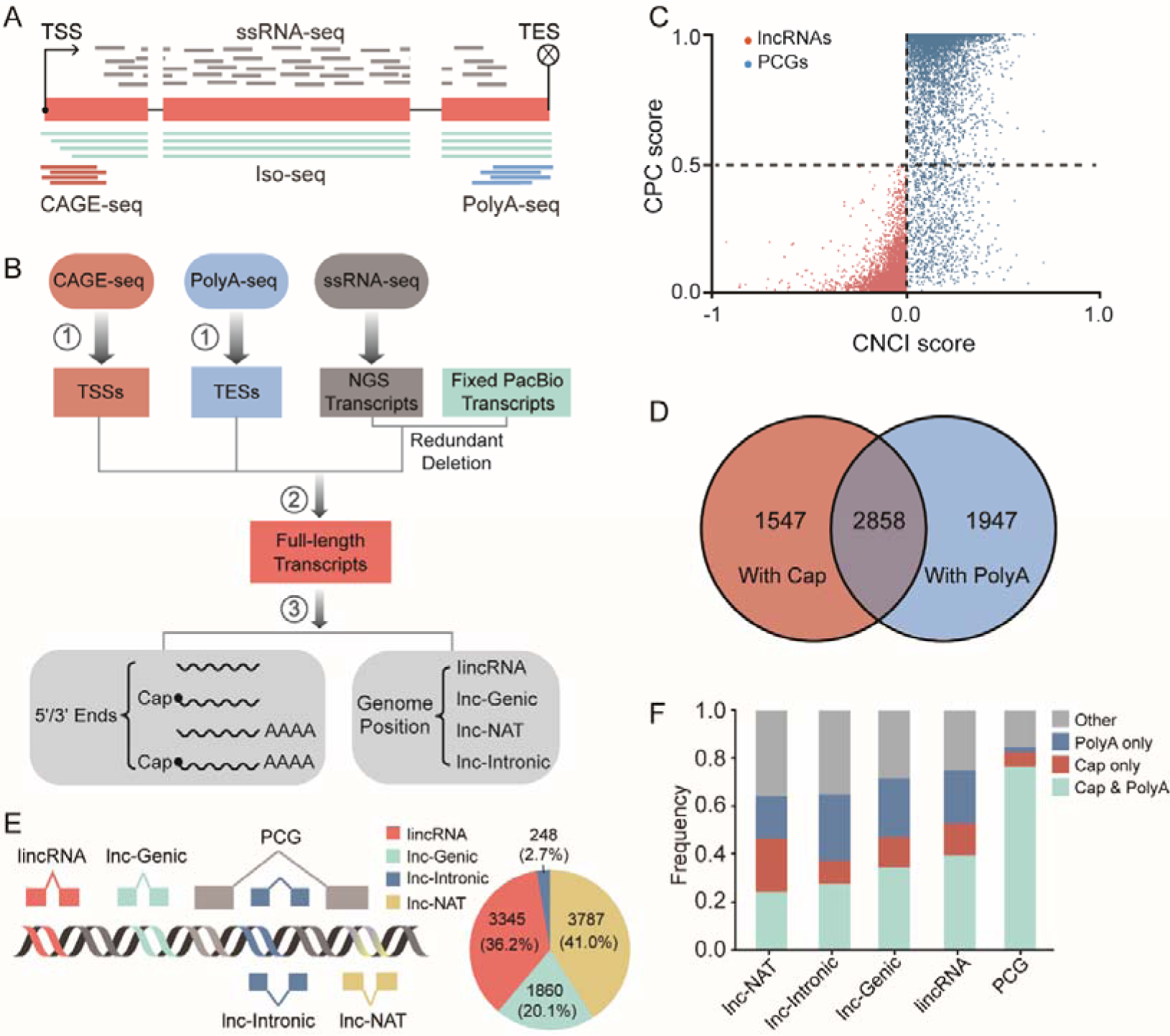
Full-length annotation of lncRNAs with multi-omics data in cotton (*G. arboreum*). **A**, The four RNA-seq technologies applied in this study. **B**, The overview of plant full-length lncRNA (PULL) pipeline. The details of procedure 1-3 were shown in Supplemental Fig. S1. **C**, Evaluation of Coding Potential Calculator (CPC) and Coding-Non-Coding Index (CNCI) for lncRNAs and PCGs. Low score indicates weak capability of protein encoding. **D**, Numbers of lncRNAs with 5’ and/or 3’ ends signals. E, Schematic illustration of four types of lncRNAs (left) and their proportion (right). F, The proportion of transcripts with 5’ cap and 3’ polyA tail in lncRNAs and PCGs.

We ultimately obtained 9240 isoforms from 8113 lncRNAs. The canonical features with CPC score < 0.5 and CNCI score < 0 are evident in the lncRNAs (**Fig. 1C**). TSSs from 4405 lncRNA transcripts and TESs from 4805 lncRNA transcripts were supported by CAGE-seq and PolyA-s eq. 2 858 lncRNA transcripts had both 5’ and 3’ end signals from CAGE-seq and PolyA-seq (**Fig. 1D**; Supplemental Table S1). According to their genomic locations, the lncRNAs could be divided into four groups: 3345 long intergenic noncoding RNAs (lincRNAs) that were located in the intergenic region (>2 Kb from PCGs); 3787 lnc-NATs that were partially or completely complementary to PCGs; 1860 long genic noncoding RNAs (lnc-Genics) (< 2 Kb from PCGs) and 248 long intron noncoding RNAs (lnc-Intronics) which were located in the introns of PCGs (**Fig. 1E**; Supplemental Table S1). Over 70 % of lnc-Genic and lincRNA have CAGE-seq or PolyA-seq signals or both, while lnc-NAT and lnc-Intronic are slightly fewer (> 60 %) (**Fig. 1F**).

To assess the accuracy of 5’ and 3’ end signals of lncRNAs from PULL on the basis of CAGE-seq and PolyA-seq, 20 lncRNAs for each of 5’ and 3’ end of lncRNAs were randomly selected to perform validation by employing 5’ and 3’ rapid amplification of cDNA ends (RACE). The Sanger sequencing results clearly showed that the 5’ and 3’ ends predicted by PULL were of extremely high accuracy, with a positive ratio of 95 % (Supplemental Fig. S2, S3). We found that numerous TSSs and TESs predicted by Stringtie had significant deviations from those predicted by the PULL pipeline (Supplemental Fig. S2, S3), because the Stringtie can only use ssRNA-seq data to assemble transcripts.

These above data reflect that at least a half of the lncRNAs in *G. arboreum* are similar to PCGs in the transcriptional processing of 5’ or 3’ ends. In summary, we constructed a landscape of lncRNAs in cotton *G. arboreum* and identified their accurate transcription regions in the genome.

### Structural and genomic features of lncRNAs in *G. arboreum*

LncRNAs were reported to have distinct genomic features with PCGs in *Arabidopsis thaliana* (Liu et al., 2012), maize (Li et al., 2014b) and rice (Zhang et al., 2014). The features in *G. arboreum* were also investigated. We found that cotton lncRNAs contain fewer exons than PCGs (mean value: 2.04 *vs* 5.05), and about 55 % of lncRNAs contain only one exon (Supplemental Fig. S4A). However, lncRNAs have relatively longer exons than PCGs (Supplemental Fig. S4C). The median length of lncRNAs transcript is 903 nt, while that of PCGs is 2998 nt (Supplemental Fig. S4E). Almost 90 % of lncRNAs only have one isoform (Supplemental Fig. S4G), showing that RNA transcription heterogeneity is much lower in lncRNA than in PCGs. In addition, lncRNA gene bodies have lower GC contents than PCGs (*P* value < 0.001, Mann-Whitney U-Test) (Supplemental Fig. S4I). Furthermore, we calculated lncRNA and PCG expression levels with FPKM (Supplemental Table S2, S3). The cumulative curve of expression values across 21 samples indicated that lncRNAs are expressed at lower levels than PCGs (Supplemental Fig. S4J). Then we examined these features of the different types of lncRNAs respectively. We found that lnc-NATs also contain shorter, but more exons, longer transcripts, lower expression levels, and higher GC content (Supplemental Fig. S4), which is different from the features of other types of lncRNAs. The features of lncRNAs in *G. arboreum* have a lot of similarities to those in other plant species (Liu et al., 2012)(Li et al., 2014b)(Zhang et al., 2014)(Wang et al., 2015a). It shows again that lncRNAs in plants share similar features in terms of genome structure and expression, although lncRNAs are modestly conserved in sequence across plant species (Deng et al., 2018).

Previous study showed that TE is highly enriched in the *G. arboreum* genome, reaching up to 68 % (Wang et al., 2016). Herein, we found that 48 % of lncRNAs contain TE-derived sequences (overlap with TEs by at least 10 nt) (Supplemental Fig. S5A). LncRNAs with more than half of TE sequences account for almost a quarter of all lncRNAs (2125 out of 9240, 23 %) (Supplemental Fig. S5A). The lincRNAs have the highest ratio of overlapping with TE (62 %), followed by lnc-Intronics, lnc-Genics, and lnc-NATs (Supplemental Fig. S5B). Besides, by comparing TE composition in lncRNAs and PCGs, we found that the gene body regions of PCGs, including 5’ UTRs, 3’ UTRs, exonic-CDS, and introns contain more DNA transposons (Class II TEs, such as *Helitron* and *TIR*) than the whole genome. While the lincRNAs (lncRNAs from intergenic regions) have the lowest composition of DNA transposons, but possess the highest composition of *Gypsy*, a kind of retrotransposon (Class I TEs) (Supplemental Fig. S5C).

We further investigated the flanking sequence of the transcription region of lincRNAs. Compared to PCGs, the nearby regions in lincRNAs also have higher sequence compositions for *Gypsy* (Supplemental Fig. S5D). The closer physical distance indicates that *Gypsy* may have a higher possibility to affect or regulate lincRNAs’ transcription. In line with our expectation, the higher expression correlation of lincRNA to nearest *Gypsy* as compared to that of PCG to its nearest *Gypsy* was observed (Supplemental Fig. S5E). These results indicate that TEs might play roles in sequence composition of lncRNAs, and suggest a possible hypothesis that TEs involvethe transcriptional expression of lncRNAs.

### *Cis* regulation of lncRNAs on expression of adjacent PCGs

Previous studies reported that lncRNAs tend to exhibit spatiotemporal expression specificity across tissues in plants (Golicz et al., 2018). To investigate whether it is the same in *G. arboreum*, we analyzed the specificity of lncRNAs expression in different tissues (details in methods). **Figure 2A** shows that the four types of lncRNAs have obvious tissue specificity, with the degree of specificity ranking from high to low as follows: lnc-NAT > lincRNA/lnc-Intronic/lnc-Genic > PCG.

**Figure 2.**
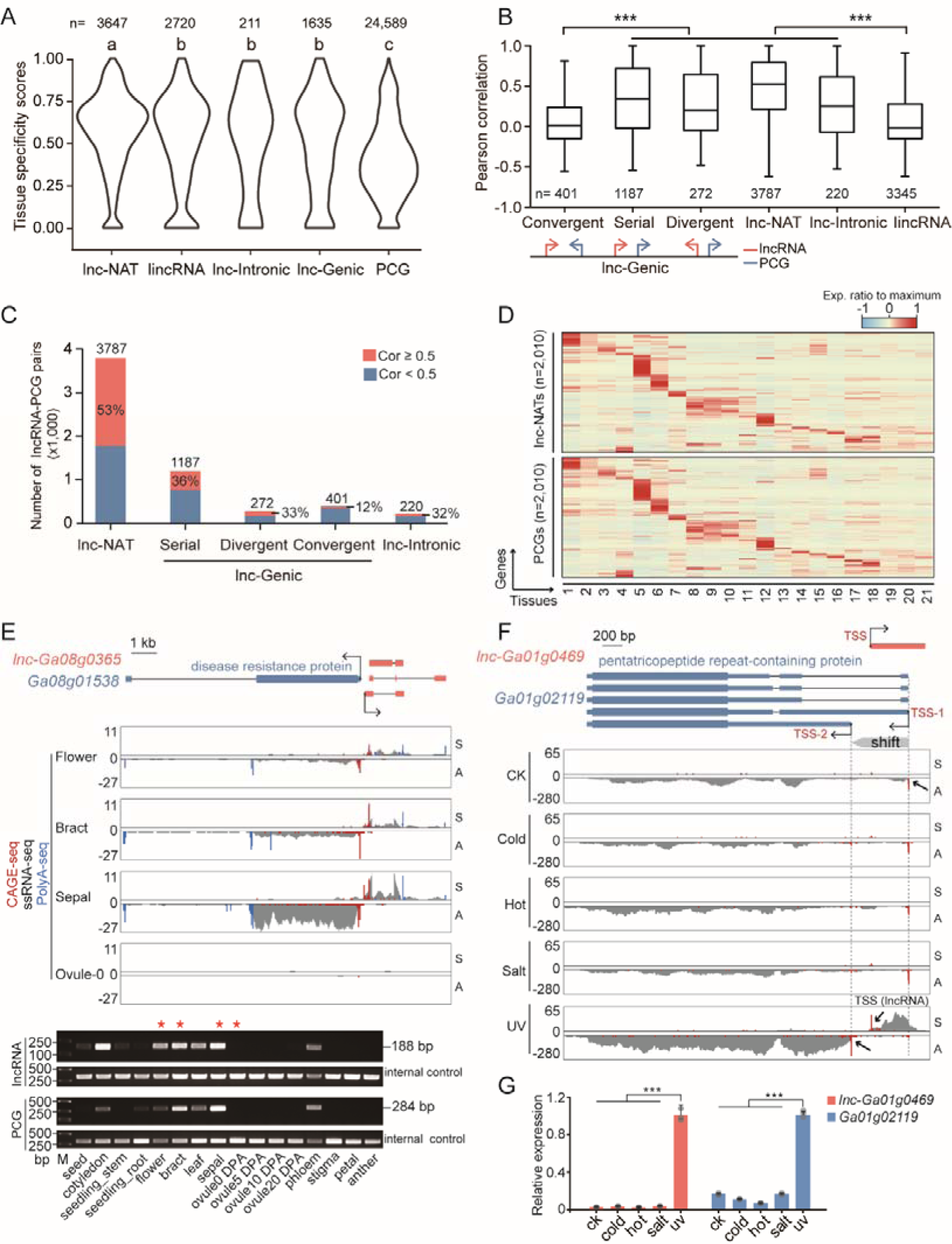
LncRNA regulates expression of adjacent PCG in *cis*. **A**, The violin plot showing the tissue specificity scores (the maximum value ‘1’ indicates a perfect tissue-specific pattern) of lncRNAs and PCGs. The scores were calculated by Jensen– Shannon divergence with ssRNA-seq data across 16 tissues. The one-way ANOVA analysis and Tukey honest significant difference post hoc test were used. **B**, Expression correlation between lncRNAs and their closest PCGs. The significance levels were indicated by asterisks (Mann-Whitney test, *** *P* value < 0.001). **C**, The number of co-expressions of lncRNA-PCG pairs. The percentage of the pairs with correlation coefficients > 0.5 (red) were shown. **D**, Heat maps showing the concordant expression of 2010 lnc-NATs (top) and their complementary PCGs (bottom) across 21 samples. The rows in top panel were arranged based on k-means clustering of lnc-NATs, while the rows in bottom panel for complementary PCGs were in the same order. Their expression values (FPKM) were scaled by z-score in row direction. **E**, The example of a lnc-Genic co-expressing with its closest PCG. S and A represent the sense and antisense strands, respectively. Semi-quantitative RT-PCR (semi-qRT-PCR) validation assays are shown below. Red stars indicate the four tissues with RNA-seq data shown above. **F**, A lncRNA related TSS switching of a PCG under UV stress. TSS-1 and TSS-2 represent the two switching TSSs. S and A represent the sense and antisense strands, respectively. **G**, The qRT-PCR validation of the up-regulation of lncRNA and PCG under UV treatment. The significance levels are indicated by asterisks (two-tailed t-test, error bar represents standard deviation, *** *P* value < 0.001).

Emerging evidence supports that some lncRNA loci act locally (in *cis*) to regulate the expression of nearby PCGs (Engreitz et al., 2016). To investigate *cis* regulation of lncRNAs in *G. arboreum*, we examined the expression correlation between lncRNAs and their nearby PCGs across tissues. The results showed that serial and divergent lnc-Genic, lnc-NAT and lnc-Intronic types exhibited significant expression correlation with their surrounding PCGs (**Fig. 2B**), while the exception is the convergent-type of lnc-Genic which forms a tail-tail type with PCGs.

Notably, expression correlation between lnc-NATs and complementary PCGs was the highest. Over half (53 %) had Cor ≥ 0.5 (**Fig. 2C**), suggesting that lnc-NATs have a strong tendency to express synchronously with their complementary PCGs. The concordant expression patterns of 2,010 lnc-NATs and their complementary PCGs (Cor ≥ 0.5) across 21 tissues are shown in **Fig. 2D**. Supplemental Fig. S6 shows a representative example of lnc-NAT in which ssRNA-seq, CAGE-seq, PolyA-seq and RT-PCR consistently supported the co-expression relationships across tissues.

In addition, several serial and divergent types of lnc-Genic identified in this study also showed a high expression correlation with adjacent PCGs (36 % and 33 % of their Cor ≥ 0.50) (**Fig. 2C**). Considering their genomic location, the two types of lncRNAs might resemble the promoter-associated lncRNA upstream antisense (ua) RNAs, promoter upstream transcripts (PROMPTs) and enhancer lncRNAs reported in mammals (Wu et al., 2017). **Figure 2E** shows a divergent type of lnc-Genic, *lnc-Ga08g0365*, in which expression is highly synchronized with adjacent coding genes across tissues.

Intriguingly, we found a phenomenon of gene transcriptional regulation which might be mediated by lncRNAs. (**Fig. 2F**). Under UV treatment, a PPR protein-encoding gene, *Ga01g02119*, switched the distal TSS (TSS-1) to proximal TSS (TSS-2), accompanied by transcriptional initiation of a lncRNA *lnc-Ga01g0469* on the antisense strand of the TSS switch region. The qRT-PCR supported the UV-induced expression of *lnc-Ga01g0469* and *PPR* (*Ga01g02119*) (**Fig. 2G**). Totally, 64 such examples were found in the genome (Supplemental Table S5). It will be extremely interesting to check whether lncRNAs could cause TSS switches of PCGs. These results indicate that lncRNAs might have diverse correlations with adjacent PCGs, including the possibility of affecting the expression intensity and tissue specificity, as well as the promoter selection of PCGs.

### TSS and TES switches of lncRNAs

RNA transcription in eukaryotes relies on three evolutionarily conserved RNA polymerases, Pol I, II, and III. Plants have evolved other two RNA polymerases, Pol IV and V (Liu et al., 2015). The studies in animals have shown that most lncRNAs are derived from transcription by Pol II, carrying a 5’ cap and 3’ polyA tail (Wu et al., 2017). However, current studies in plants still lack enough evidences. The accurate definition of 5’ and 3’ ends for lncRNA transcripts in cotton gives us the clue to address the issue.

Firstly, we performed motif discovery for lncRNA TSSs and TESs using Homer software (Heinz et al., 2010). As shown in Supplemental Fig. S7, the TATA and Y (pyrimidine) patch of promoter (Yamamoto et al., 2009), and PAS and U-rich of terminators (Wang et al., 2018a) for a canonical Pol II transcription have been identified around TSSs/TESs of lncRNAs. Next, we surveyed the nucleotide composition of ±50 nts of TSSs and TESs of lncRNAs and found that nucleotide composition around TSSs and TESs has similar bias with that of PCGs (**Fig. 3, A and C**; Supplemental Fig. S8), resembling TSSs of PCGs in *Arabidopsis* (Tokizawa et al., 2017) and TESs of PCGs in rice (Fu et al., 2016). We also analyzed the polymerase II (Pol II) chromatin immunoprecipitation sequencing (ChIP-seq) data of leaves according to previous method (Zhao et al., 2018a). 3390 high-confidence binding peaks were identified in the *G. arboreum* genome. The similar ratio (13 % vs 10 %) mapping to the expressed lncRNAs and PCGs in leaves (FPKM >= 1). These results support that most of lncRNAs in cotton are transcribed by Pol II.

**Figure 3.**
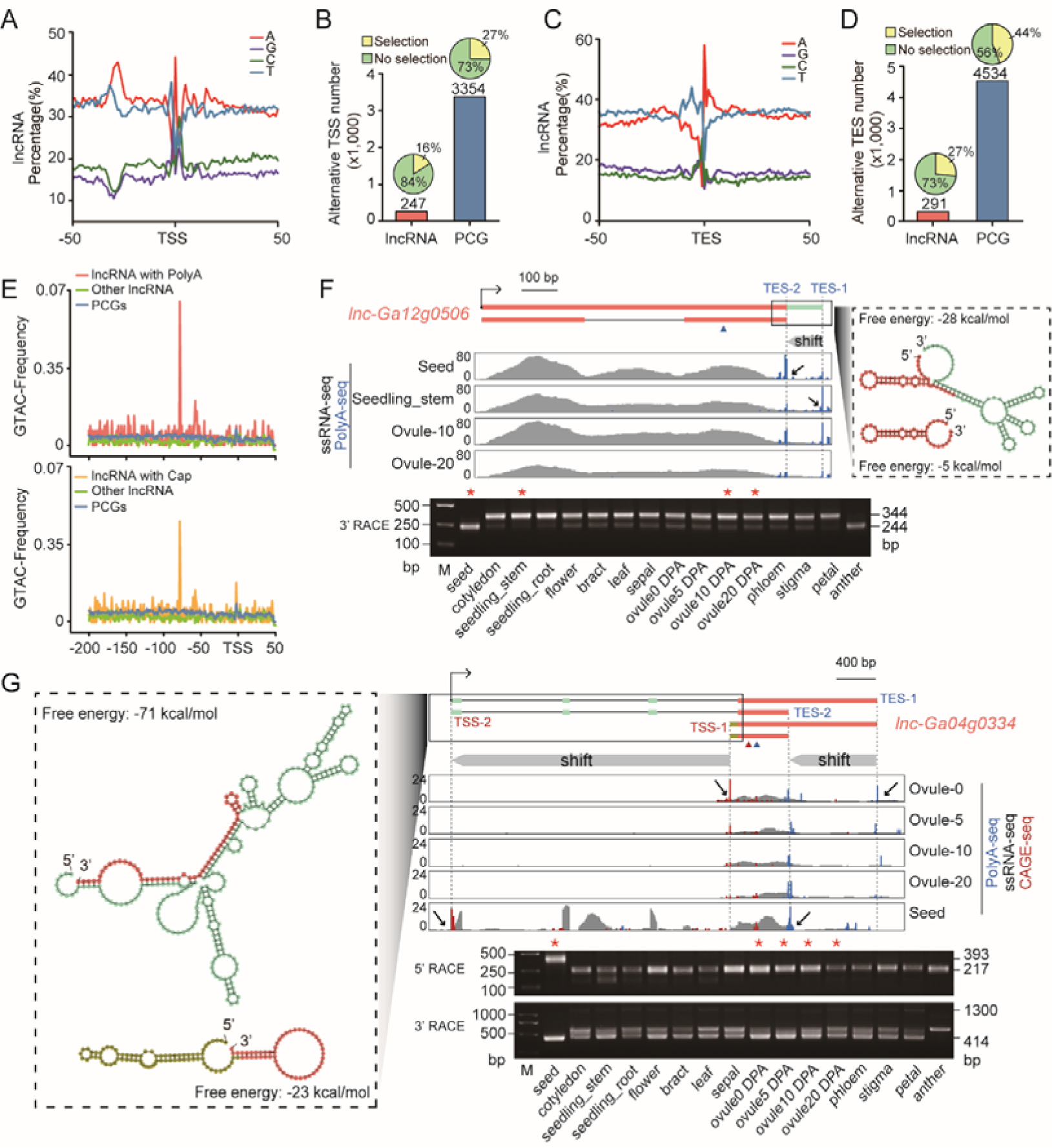
Alternative usage of TSSs and TESs of lncRNAs. **A**, Nucleotide composition at ±50 nt around TSSs in lncRNAs. **B**, Numbers of multi-TSSs in lncRNAs and PCGs; Pie charts indicate the percentage of TSS switching across tissues for lncRNAs and PCGs. **C**, Nucleotide composition at ±50 nt around TESs in lncRNAs. **D**, Numbers of multi-TESs in lncRNAs and PCGs; pie charts indicate the percentage of TES switching across tissues for lncRNAs and PCGs. **E**, A DNA motif was identified in promoter regions of lncRNAs with 5’ cap or 3’ polyA. Other lncRNA represents lncRNAs without 5’ cap and without 3’ polyA. **F**, An example of a lncRNA with alternative TES usage across tissues. PolyA-seq and ssRNA-seq signals (Top left), RNA secondary structure (Top right), and validation with 3’ RACE assays (Bottom) are shown. TES-1 and TES-2 represent the two switching TESs. The blue triangle indicates the position of RT primer for the 3’ RACE. Red stars indicate the four tissues with RNA-seq data shown above. **G**, The lncRNAs with alternative TSS and TES usage across tissues. The CAGE-seq, PolyA-seq and ssRNA-seq signals (Top right), RNA secondary structure (Left), and validation with 5’ and 3’ RACE assays (Bottom right) were shown. TSS-1 and TSS-2, TES-1 and TES-2 represent the two switching TSSs and TESs, respectively. The red and blue triangles indicate the positions of RT primers for the 5’ RACE and 3’ RACE, respectively. Red stars indicate the five tissues with RNA-seq data shown above.

Above results support that most of lncRNAs in cotton are transcribed by Pol II and processed with 5’ m7G caps and 3’ polyA tails. Notably, an unknown 4-nt motif, GTAG, was found to be significantly enriched around 75 nt upstream of the TSSs (**Fig. 3E**) in some lncRNAs with 5’ caps or 3’ polyA tails. The function of this motif for lncRNA transcription deserves in-depth study. However, for lncRNAs without the 5’ and 3’ end signals, these features of Pol II transcription were not detected (Supplemental Fig. S9), indicating that transcription of those lncRNAs (∼25%) in cotton might depend on other RNA polymerases.

To date, the genome-wide investigation on TSS/TES selection regulation for lncRNA was rare. Herein, we performed the pilot investigation in *G. arboreum*. Using the CAGEr program and requiring TPM >0.5 in every tissue, we determined 3,857 TSSs and 4279 TESs corresponding to 3709 and 4067 lncRNA loci, respectively. There were in average 1.04 TSSs and 1.05 TESs per lncRNA, which was lower than PCGs. 6.7% and 7.2% of lncRNAs are multi-promoter and multi-terminator (≥2 TSSs/TESs) (Supplemental Fig. S8, C and D) RNAs. Next, we performed comparative analysis on TSS/TES usage dynamics across 21 tissues and identified 40 and 77 switching events for TSSs and TESs of lncRNAs between any two tissue samples. This accounted for 16.2% and 26.7% of multi-promoter and multi-terminator lncRNAs, respectively (**Fig. 3, B and D**).

Although alternative use of TSS/TES was lower in number and frequency in lncRNAs than that of PCGs (**Fig. 3, B and D**), we suggest that it still deserves attention. For *lnc-Ga12g0*506, differential usage of TES sites across tissues would cause significant 3’ end length and structure changes (**Fig. 3F**). The *lnc-Ga04g0334* has two TSSs and two TESs during ovule development. With the development from young ovules to matured seeds, switching of both the TSS and TES of the lncRNA occurs (**Fig. 3G**). In the seeds, a longer transcript completely changes its original 5’ terminal secondary structure. Other examples of lncRNAs with TSS/TES switches are shown in Supplemental Fig. S8, E and F. Because the function of lncRNAs includes regulating chromatin topology and scaffolding protein or RNAs (Ransohoff et al., 2017), changes in length produced by TSSs/TESs switch will inevitably alter the structure of lncRNA, thereby possibly change its function.

### Genetic variations and GWAS sites associated lncRNAs of *G. arboreum*

As PCGs are subject to strong selection pressure to conserve protein sequences during evolution, their coding sequences maintain a lower mutation frequency. To investigate the genetic selection pressure in lncRNAs, statistical analysis for natural mutation frequency distribution of lncRNA locus was performed. The 17,883,108 high-quality SNPs were derived from natural mutations in 230 *G. arboreum* lines (Du et al., 2018). As expected, the mutation frequency of lncRNAs was higher than that of PCGs overall. However, similar to PCGs, mutation frequency in lncRNA gene bodies (2.85 SNP/kb) was significantly lower than its flanking sequence, and the mutation frequency of TSS upstream was significantly lower than TES downstream (3.19 vs 3.91 SNP/kb). Exons and introns, and their partial flanking sequences also revealed a similar trend as PCGs in mutation frequency curves (**Fig. 4A**). An intriguing phenomenon worth noting, is that the mutation frequency in lncRNA gene bodies is the same as the TSS upstream region of PCGs (**Fig. 4A**). To rule out the possible influence of sequence overlap between PCGs and lncRNAs, we only used the lincRNAs to perform the analysis, and the same trend was observed (Supplemental Fig. S10).

**Figure 4.**
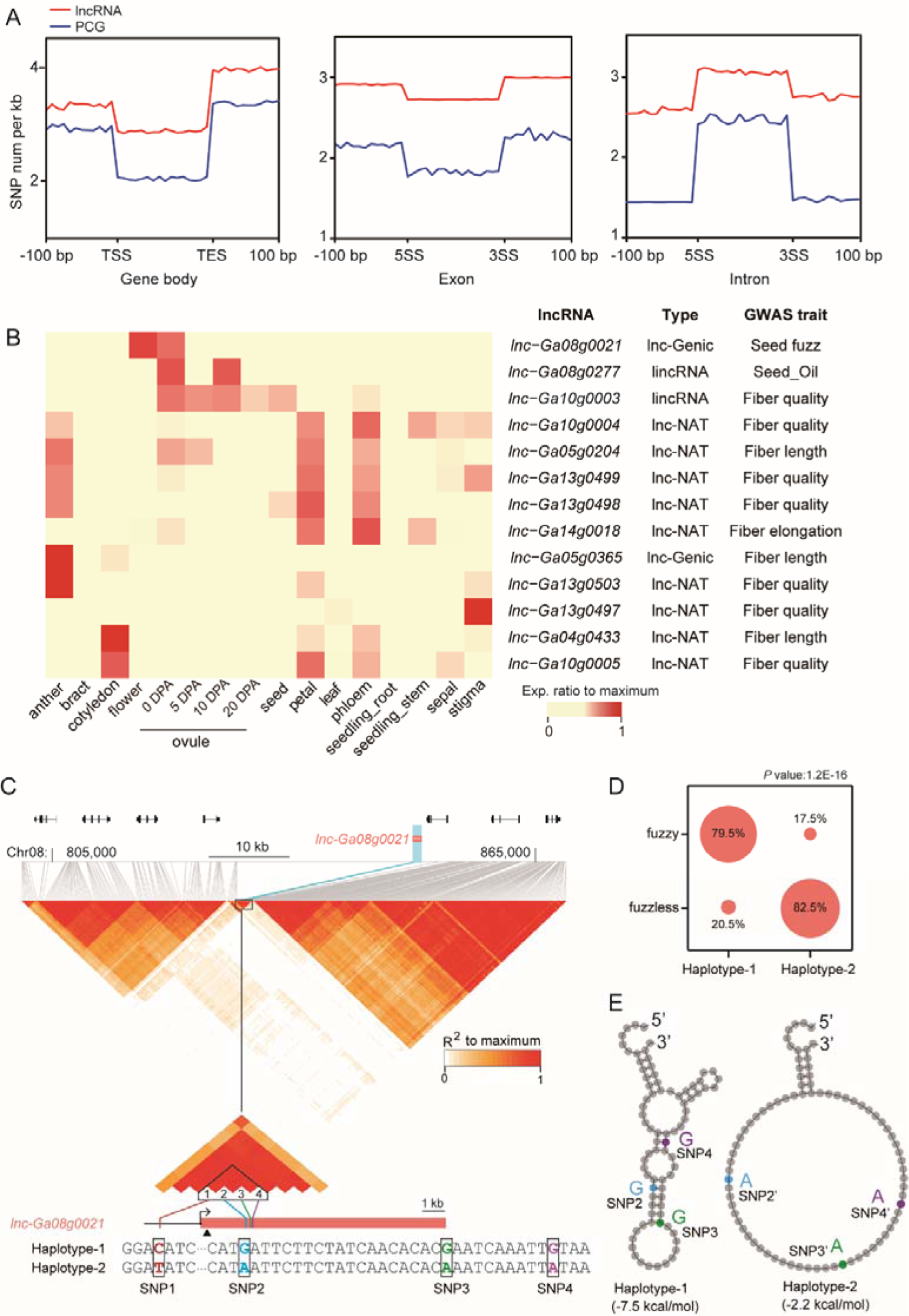
Genetic variations and linkage to GWAS of lncRNAs in ovule development. **A**, SNP frequency distribution on the gene body (Left), exon (Middle) and intron (Right) of lncRNAs (Red lines) and PCGs (Blue lines). **B**, The lncRNAs linked to GWAS traits. The heatmap (Left) shows tissue expression of the lncRNAs. **C**, The linkage disequilibrium (LD) plot of *lnc-Ga08g0021* locus. The zoom-in LD plot and the sequence of two haplotypes were shown in bottom. One SNP in promoter and three SNPs in gene body of *lnc-Ga08g0021* were highlighted. The black triangle indicated the target site of gra-miR8739. **D**, The bubble chart indicating the seed phenotype of the two haplotypes. The *P* value was marked above (Chi-squared test). **E**, The RNA secondary structures of two haplotypes. The minimum free energy was given below.

According to previous studies (Mercer et al., 2009), lncRNAs can recruit chromatin remodeling or mediate DNA and histone modification to achieve effects on PCG promoters or enhancers. LncRNAs and their target DNA elements in promoters are directly or indirectly combined together to form transcriptional regulatory machinery in the regulatory region of PCGs. Therefore, the sequence mutation from the lncRNA or promoters of PCGs would affect their downstream gene expression, suggesting that they are under the same genetic selection pressure, and experience comparable mutation frequencies.

In addition, GWAS sites associated with cotton agronomic traits have been identified previously (Fang et al., 2017)(Wang et al., 2017)(Du et al., 2018)(Hou et al., 2018)(Li et al., 2018)(Ma et al., 2018). To discover the possible links between lncRNAs and cotton agronomic traits, we mapped the 932 GWAS sites into lncRNAs loci and found the GWAS sites overlapping with the gene body regions of 13 lncRNAs, including two lincRNAs, nine lnc-NATs and two lnc-Genics. The tissue expression analysis showed that three lncRNAs: one lnc-Genics and two lincRNAs exhibited obvious tissue-specific expression during ovule development (0-10 DPA). The three GWAS sites associated with seed fuzz (*lnc-Ga08g0021*), seed oil accumulation (*lnc-Ga08g0277*) and fiber quality (*lnc-Ga10g0003*), respectively (**Fig. 4B**). We then further analyzed the GWAS region controlling seed fuzz based on previous study (Du et al., 2018). As shown in **Fig. 4C**, a distinct linkage disequilibrium (LD) block encompassing the *lnc-Ga08g0021* and a PCG overlaps with the peak signal of GWAS. Based on the four SNPs in the block across 230 *G. arboreum* lines, the *lnc-Ga08g0021* could be divided into two haplotypes (**Fig. 4C**) which were closely related to fuzz and fuzzless phenotypes, respectively (**Fig. 4D**). The RNA structure analysis showed that the three SNPs in transcript of the lncRNA could lead to great structural variation (**Fig. 4E**). Intriguingly, we found that the *gra-miR8739* (a miRNA in *G. raimondii*) could target this lncRNA (Supplementary Table 1). Thus it could be a candidate trans-acting siRNA producing locus (TAS) with potential to produce ta-siRNAs or phased siRNAs. So we infer that the RNA structural variation may affect the ta-siRNA processing, and finally cause the different seed phenotypes. These results reveal that lncRNAs might be involved in the developmental regulation of ovules and fiber, either alone or with their neighboring PCG.

### Regulatory functions of lncRNAs in *G. arboreum*

To reveal the puptative functions of lncRNAs, we applied the weighted correlation network analysis (WGCNA) to analyze the co-expression between lncRNAs and PCGs by using transcriptomic data from 21 samples. A total of 7684 lncRNAs and 25,253 PCGs were integrated in the network construction after filtering genes with low expression (FPKM < 10 for PCGs, FPKM < 1.5 for lncRNAs). Finally, 43 co-expressing modules were identified (**Fig. 5A**; Supplemental Table S6). Modules 16 and 13 contain the homologous genes of *GhMYB25-like* and *GhEX1* in *G. hirsutum*, which were previously identified as key regulators of fiber initiation and elongation (Wang et al., 2016)(Shan et al., 2014). The two lincRNAs in these two module were selected to test their expression by qRT-PCR. The results reflect that both of the lncRNAs and PCGs exhibit ovule-specific expression (**Fig. 5B**).

**Figure 5.**
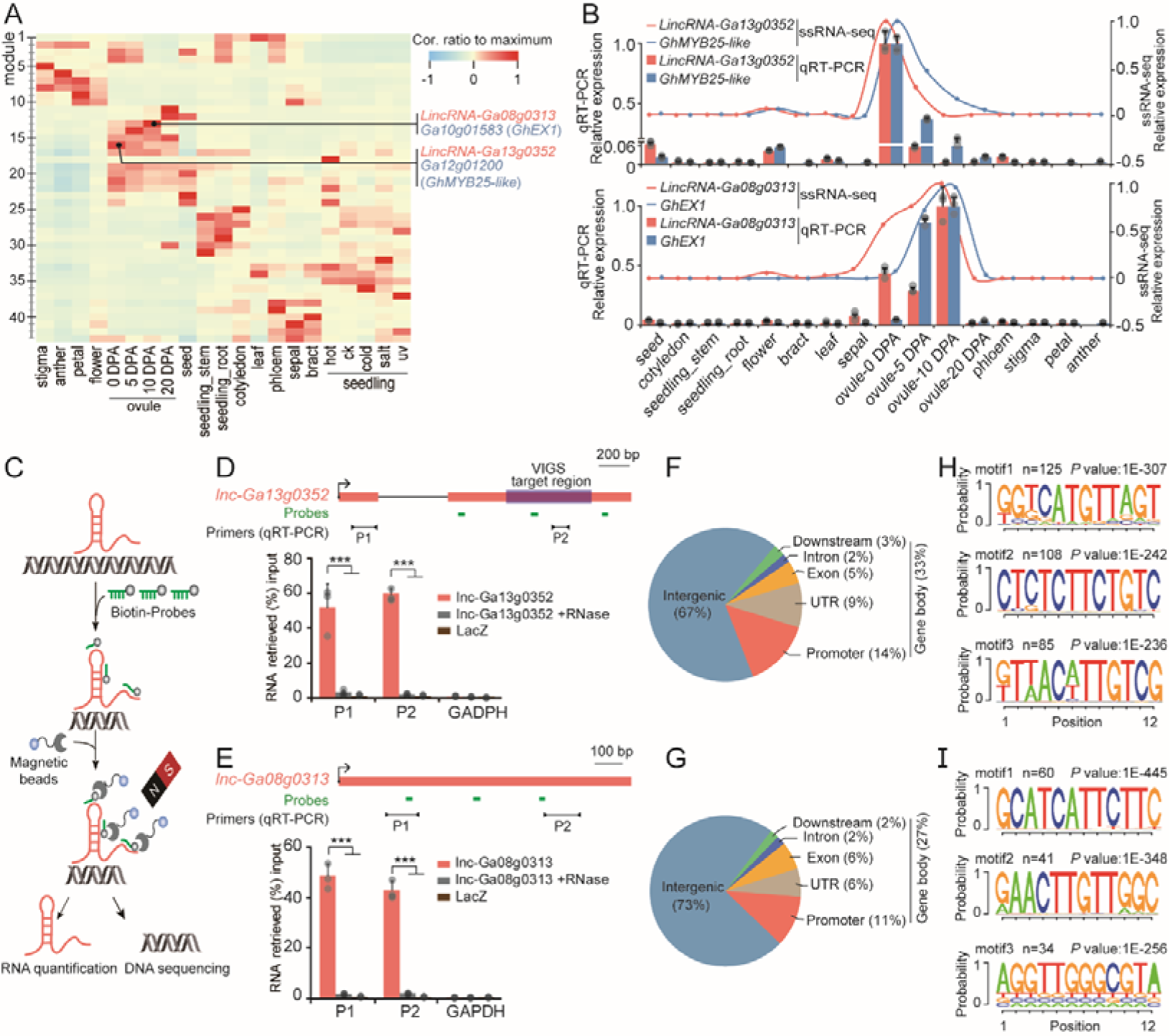
ChIRP-seq analysis reveals genome-wide binding sites for *lnc-Ga13g0352* and *lnc-Ga08g0313*. **A**, The WGCNA association analysis for the module and tissue. The modules containing known PCGs involved in cotton fiber initiation and elongation are indicated. The rows and columns correspond to modules and tissues. Each cell is color-coded by correlation according to the color legend. The color scale means the correlation coefficient (−1 to 1) between the module and tissue sample. **B**, The co-expression of lncRNA and PCG in ssRNA-seq (lines) and corresponding validation by qRT-PCR (bars), two-tailed t-test, and error bar represents standard deviation. **C**, Experimental flow chart of ChIRP. The biotinylated probes (green) for lncRNA were used. The enriched RNAs were quantified by qRT-PCR, and the enriched chromatin DNAs were used to perform DNA sequencing. **D-E**, Schematic genomic structure of two lincRNAs and the enrichment of lncRNA transcript in ChIRP. The positions of the biotinylated ChIRP probes were shown with green lines. VIGS target region of *lnc-Ga13g0352* was highlighted with purple box. P1 and P2 which were marked by black lines represent two pairs of qRT-PCR primers. The lacZ probes and RNase-treated samples were used as non-targeting control and negative control, respectively. The *GAPDH* transcript was used as a negative internal gene control. The significance levels are indicated by asterisks (two-tailed t-test, three biological replicates, error bars represent standard deviation, *** *P* value <0.001). **F-G**, Percentage of the lncRNA binding sites localization in the genome for *lnc-Ga13g0352* (F) and *lnc-Ga08g0313* (G). **H-I**, The putative three binding motifs of the *lnc-Ga13g0352* (H) and *lnc-Ga08g0313* (I).

Based on our analysis, the two lincRNAs were ruled out the possibility of pri-miRNA, ceRNA or precursor of phased siRNA (Supplemental Table S1). So we examined their putative interaction with chromatin by performing chromatin isolation by RNA purification sequencing (ChIRP-seq) (**Fig. 5C**)(Percharde et al., 2018)(Zhao et al., 2018b). The RNA enrichment detection validated by qRT-PCR showed that the biotinylated probes for *lnc-Ga13g0352* and *lnc-Ga08g0313* efficiently captured the corresponding lncRNAs (**Fig. 5, D and E**). Respectively, 969 and 440 high-credibility binding peaks in two biological replicates for the two lncRNA were found (Supplemental Fig. S11). Genome-wide distribution analysis shows that 33% and 27% of their binding peaks are in gene body region, corresponding to 317 and 114 PCGs, respectively (**Fig. 5, F and G**; Supplemental Table S7). The motif analysis of these binding sites identified three core motifs for these lncRNAs, suggesting that the two lncRNAs might bind targets in the specific DNA motifs (**Fig. 5, H and I**).

The high-enriched binding peaks of *lnc-Ga13g0352* and *lnc-Ga08g0313* were chosen to perform ChIRP-qPCR validations. The peak regions and other regions of target genes were chosen as the detection sites and negative control for qPCR, respectively (**Fig. 6, B and C**; Supplemental Fig. S12B and S13B). The significant enrichment between lncRNA-bound chromatin fragments (probes) and total chromatin input (none-probes) were observed in the examples, showing a high-credibility of our ChIRP-seq data. The GO enrichment (**Fig. 6A**; Supplemental Fig. S13 and Supplemental Table S7) analysis for the candidate PCGs regulated by the lncRNA showed that the target PCGs (ChIRP-seq peaks <3 Kb around gene body) of *lnc-Ga13g0352* displayed remarkable enrichment of transcription regulator activity (GO: 0140110, *p*-value = 1.63 x 10^-4^) and DNA-binding transcription factor activity (GO: 0003700, *p*-value = 1.51 x 10^-4^), indicating that this lncRNA might regulate genes with transcriptional functions.

**Figure 6.**
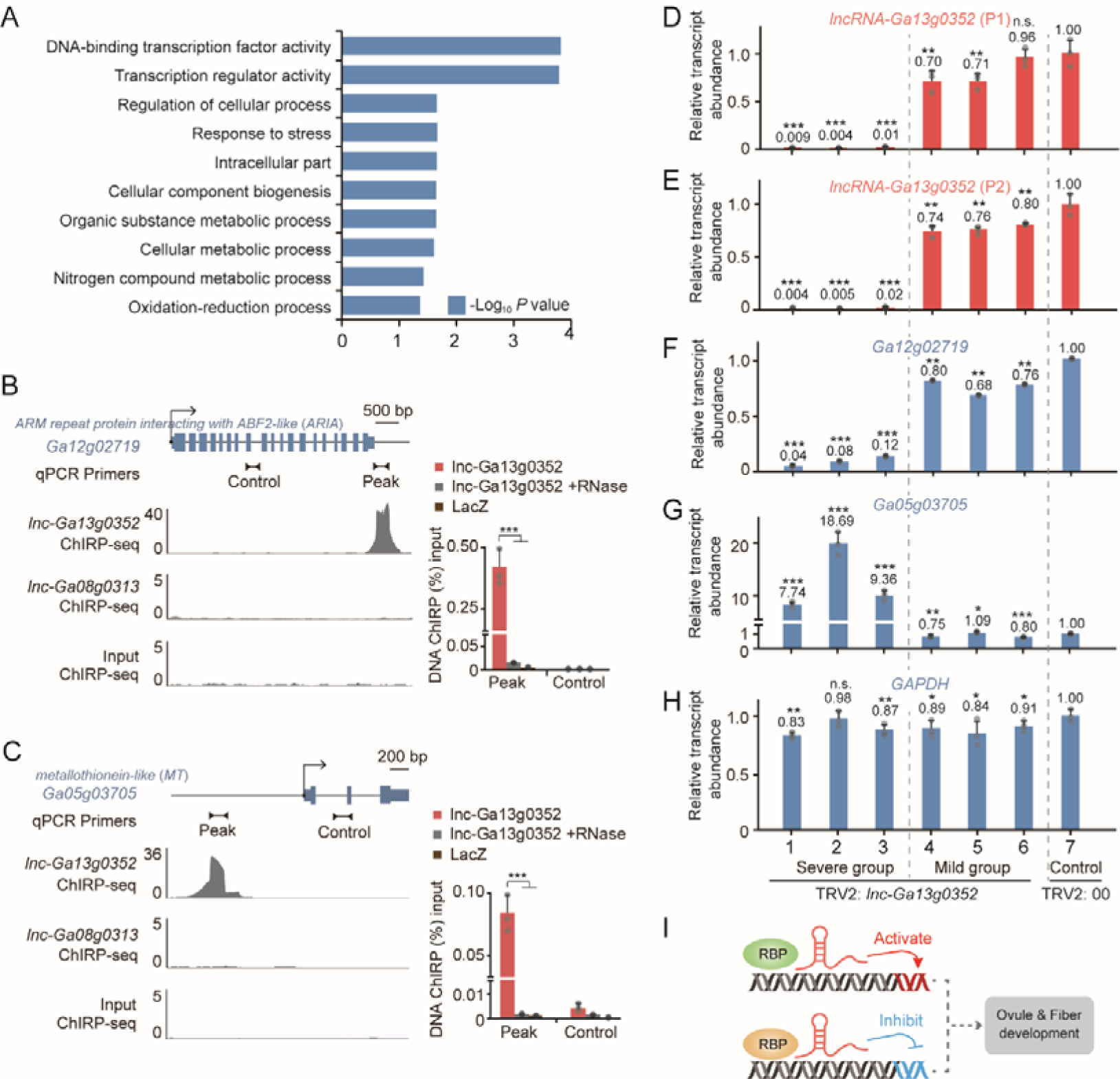
*Lnc-Ga13g0352* might regulate transcription of ovule development associated genes via activating and inhibiting manners. **A**, Top 10 GO terms in ontology enrichment for lncRNA binding genes. **B-C**, The ChIRP-seq peaks and ChIRP-qPCR for *Ga12g02719* (*ARIA*) and *Ga05g03705* (*MT*). The peaks from one biological replicate were shown. The lacZ and *lnc-Ga08g0313* probes were used as non-targeting control, and RNase-treated samples were used as negative control for ChIRP. The significance levels of qPCR are indicated by asterisks (two-tailed t-test, three biological replicates, error bars represent standard deviation, *** *P* value < 0.001). **D-G**, Relative transcription levels of *lnc-Ga13g0352* (D, E), *ARIA* (F), and *MT* (G) in the leaves of six independent VIGS seedlings (TRV2: *lnc-Ga13g0352*) and one vector control (TRV2: 00) plant. The P1 and P2 primers in Figure 6d were used for qRT-PCR of *lnc-Ga13g0352.* The significance levels are indicated by asterisks (two-tailed t-test, error bars represent standard deviation, * *P* value < 0.05;** *P* value < 0.01; *** *P* value < 0.001; n.s. represents not significant). **H**, *GAPDH* was used as a negative control for VIGS. The significance levels are indicated by asterisks (two-tailed t-test, error bar represents standard deviation, * *P* value <0.05; ** *P* value < 0.01; *** *P* value < 0.001; n.s. represents not significant). **I**, The genome-wide regulatory model of *lnc-Ga13g0352*.

Next, we wondered what the regulatory effects of the *lnc-Ga13g0352* on its targets are. We analyzed the expression pattern of 317 target PCGs across 21 tissues (Supplemental Fig. S12A). Three clusters were classified by k-means clustering (Euclidean distance metric), two of which exhibited significant differences based on their expression in ovules. 115 of the target PCGs were highly expressed in ovules (EO cluster), and 121 of the target PCGs were completely inhibited in ovules (IO cluster). Hence, we supposed that this lncRNA might regulate ovule development associated genes through two opposite manners: activation and inhibition.

Currently, there is no feasible approach for stable transformation in *G. arboreum*, so we used virus induced gene suppression (VIGS) experiments to test the effects caused by the silencing of *lnc-Ga13g0352* (**Fig. 5D**; Supplemental Fig. S12C). The ectopic silencing assay in leaves of *G. arboreum* seedlings showed that the six independent VIGS lines exhibited different silencing effects for *lnc-Ga13g0352*, and could be divided into severe group (< 5 % of control; seedling 1, 2 and 3) and mild group (∼ 75 % of control; seedling 4, 5 and 6) (**Fig. 6, D and E**). We then investigated the transcriptional expression of two target PCGs: *Ga12g02719* and *Ga05g03705* which respectively belongs to EO and IO cluster of *lnc-Ga13g0352*’s targets. The qRT-PCR results show that these two genes were inhibited and activated respectively (**Fig. 6, F-H**). Moreover, the expression of these two target PCGs exhibit dose-dependent effect in control (line 7), mild (line 4-6) and severe groups (line 1-3). The inhibition for *Ga12g02719* and activation for *Ga05g03705* in the severe group was stronger than that in the mild group. Together, above results support the genome-wide opposite regulatory roles of *lnc-Ga13g0352* for its target PCGs (**Fig. 6I**).

## DISCUSSION

Previous lncRNA identification studies in cotton only relied on RNA-seq (Wang et al., 2015b) (Zhao et al., 2018a), which would produce incomplete annotations for lncRNAs due to transcript assembly errors, especially in 5’ and 3’ ends caused by the low reads coverage, and false-positive annotation for lncRNAs caused by non-strand-specific reads or background of ssRNA-seq. In this study, we compiled PULL pipeline to integrate multi-omics data to accurately annotate lncRNAs. Compared with previous lncRNA identification in leaf tissue of *G. arboreum* (Zhao et al., 2018a), this is a more comprehensive and accurate lncRNA dataset, and it updated a considerable number of lncRNAs with accurate 5’ or 3’ ends or both. Based on these accurate termini annotation for lncRNAs supported by CAGE-seq and PolyA-seq, we found that most of lncRNAs in *G. arboreum* have 5’ caps and a polyA tails. Moreover, the canonical Pol II transcription elements were also identified in their promoters and terminators. These results provide new evidence to support that plant lncRNAs are transcribed by RNA Pol II, which was in line with previous studies (Zhao et al., 2018a).

The compositional contribution of TEs to lncRNAs in plants has been reported in previous studies (Zhao et al., 2018a)(Golicz et al., 2018). In this study, we also came to the similar conclusion that retrotransposons, especially *Gypsy,* contributes the most to the composition and transcriptional regulation of lncRNAs in cotton (*G. arboreum*). If given that different plant species experienced biased TE explosion events (Lisch, 2013) and different TEs would have different insertion preferences (Sigman and Slotkin, 2016), it seems plausible that the lncRNAs derived from TEs would exhibit tremendous differences in sequence composition and genomic location among different plant species, and even in closed species in the same lineage (Zhao et al., 2018a), which explains the fact that low conservation of lncRNAs among plant species.

Previous studies provided sporadic evidence supporting the important biological functions of alternative TSSs/TESs of lncRNAs, such as (1) alternative polyadenylation of an antisense transcript in an important *Arabidopsis* floral repressor gene *FLC*, which controls *FLC* expression and thus flowering time (Csorba et al., 2014); (2) a lncRNA named *SVALKA* in *Arabidopsis* that modulates response to cold via the acclimation gene C-repeat/dehydration-responsive element binding factor 1 (*CBF1*) through lncRNA read-through transcription (Kindgren et al., 2018). Therefore, it is foreseeable that the systematic investigation of TSS/TES switching of lncRNAs will uncover more cases that are involved in the control of development or stress response The present study has pioneered a genome-wide identification and dynamic monitoring of TSSs and TESs for lncRNAs in crops. The novel findings in this study deserve further attention.

Our results also showed that some lncRNAs might be correlated to the stress-triggered TSS switching of PCGs (**Fig. 2, F and G**), and have high tissue-specific expression and concordant expression with nearby PCGs (**Fig. 2, A and E**). Hence, the heterogeneity of the lncRNA transcription and its possible regulation on transcription of PCGs might make cardinal contributions to the rewiring of gene regulatory networks in plants. It will be of great value to address the biogenesis and mode of action of these novel transcription events.

Our ChIRP-seq explored the genome-wide binding sites of*lnc-Ga13g0352* and its VIGS assay indicated that this lncRNA might impose two distinct effects on the transcriptional expression of its target PCGs: activation and inhibition. Our study not only uncovered the potential regulation function of the lncRNA on cotton development, but also shed light on the complex mode of action for lncRNAs. How a lncRNA can modulate gene expression network in two opposite manners on genome-wide is worth further study.

## MATERIALS AND METHODS

### Plant Growth and Treatments

Cotton *G. arboreum* cv. Shixiya1 (SXY1) was cultivated in an automated greenhouse where the temperature was set to 30 ℃/25 ℃ for 16 hours day/8 hours night cycle and the humidity was set to 65%. The matured tissue samples included anther, stigma, petal, bract, sepal and whole flower at 0 days post-anthesis (DPA), and phloem, leaf, seedling root, seedling stem, cotyledon (2 weeks), seed, and ovules at four developmental stages (0, 5, 10 and 20 DPA). For stress treatment samples, seedlings (2 weeks) were used. Stress conditions included hot (50 ℃ for 5 hours), cold (5 ℃ for 15 hours), salt (500 mM NaCl) and UV (1.24 *μ*mol m^−2^ s^−1^ UV-B for 2 hours). Untreated seedlings were used as control.

### RNA-seq and PCR Assays

Cotton RNA extraction and libraries construction for ssRNA-seq, CAGE-seq and PolyA-seq were performed according to our previous publication (Wang et al., 2019b). PacBio sequencing data of mixed samples from 16 tissues was from our previous study (Wang et al., 2019b). For qRT-PCR, 1 ug total RNA were used to synthesize cDNA with oligo dT or gene-specific primers by using reverse transcription kit (Vazyme, China). Relative gene expression was calculated using the 2^-ΔCT^ method.

The internal control *Ga01g02318* in our study (Wang et al., 2019b) was used for semi-qRT-PCR and qRT-PCR. For 5’ and 3’ RACE, the nested PCR products were analyzed by Qsep100DNA Analyzer (BiOptic, Taiwan, China) and sent to TSINGK Company for Sanger sequencing. All primers used in this study are listed in Supplemental Table S8.

### Sequencing Data Processing

For ssRNA-seq data, raw reads were first processed by adapter trimming and removing low quality or inaccurate reads. Then clean reads were mapped to the recently updated reference genome of *G. arboreum* (Du et al., 2018) with STAR (Dobin et al., 2013) in EndToEnd and 2-pass mapping modes. For CAGE-seq and PolyA-seq data, raw reads were treated similarly to get clean reads. Then clean reads were mapped by STAR with EndToEnd mode. The removal of PCR duplicates was achieved by FastUniq (Xu et al., 2012). The isoforms assembled by fixed Iso-seq data were adopted from our previous study (Wang et al., 2019b). Supplemental Table S10 lists all software and associated information regarding the version and options/parameters.

### PULL Pipeline

Alignment results from ssRNA-seq and Iso-seq were merged by StringTie (Pertea et al., 2015) with parameter -g 100. The program ‘clusterBed’ in Bedtools (Quinlan and Hall, 2010) was used to get a whole non-redundant transcriptome collection. TSSs/TESs from CAGE-seq/PolyA-seq were obtained by R package, CAGEr (Haberle et al., 2015). To match a transcript with a TSS supported by CAGE-Seq, we considered two factors: (1) “dis”, the distance between a TSS from CAGE-seq and the 5’ end of the assembled transcript (Hon et al., 2017). (2) “cor”, the coefficient between transcripts per million (TPM) of a TSS from CAGE-seq and fragments per kb of exonic sequence per million mapped reads (FPKM) of the transcript from ssRNA-seq across samples (Kawaji et al., 2014). For each assembled transcript, we retained all CAGE peaks between 500 bp upstream of the 5’ end and the first exon. CAGE peak with the highest “cor” was selected as the TSS of the transcript. If there were multiple CAGE peaks with the same “cor”, the CAGE peak with the smallest “dis” was selected as the TSS of the transcript. The matching of a transcript with a TES from PolyA-seq was followed a similar way.

To get a set of lncRNA transcriptomes, the following assembled transcripts were directly removed: (1) the known PCGs annotated in the genome and known ncRNAs in Rfam (tRNAs, rRNAs, snoRNAs, and snRNAs); (2) the transcripts with lengths less than 200 bp or ORF lengths more than 100 amino acids (aa); (3) the transcripts encoding known protein in Swiss-Prot or Pfam; (4) the transcripts with Coding-Non-Coding Index (CNCI) score > 0 (Sun et al., 2013) or Coding Potential Calculator 2 (CPC2) score > 0.5 (Kang et al., 2017); Besides, for the candidate long noncoding natural antisense transcripts (lnc-NATs) without TSS or TES signals from CAGE-seq or PolyA-seq, we used the ‘intersectBed’ program to filter the contamination from sense transcripts in ssRNA-seq. If their complementary sequence length occupied more than 50 % of the length for both lnc-NATs and complementary PCGs, those lnc-NATs were removed. The specific implementation method can be referred to at the PULL pipeline at https://github.com/cotton-lab/PULL. To annotate the candidates of pri-miRNAs, precursors of phased siRNA, and competing endogenous RNAs (ceRNAs) in the lncRNAs, all the miRNAs of *Gossypium* genus (miRbase version 22.1) were used to perform prediction by RNAfold (v 2.4.14) (Lorenz et al., 2011), psRNATarget (Dai et al., 2018) and PerRBase (Yuan et al., 2017). The 8213 and 65 candidates of pri-miRNAs, precursor of phasiRNA, and ceRNAs were annotated in Table S1.

### Transposable Element (TE) Analysis

The TE annotation in GTF format was obtained from previous study (Du et al., 2018). The FPKM values of TE were calculated using StringTie. Considering that many TEs are repetitive, multi-mapping reads were dropped, only unique mapping reads being used to calculate FPKM. The lncRNAs or PCGs containing TE sequences with the lengths of more than 10 bp were considered as TE-related lncRNAs or PCGs, which were achieved by ‘intersectBed’ program in Bedtools. Pearson correlation coefficient was calculated between the expression levels of the TE and TE-related lncRNA or PCG across samples. When comparing the correlation between TE-lncRNA or TE-PCG pairs, we only chose the pairs without overlap to avoid its impact on FPKM values.

### Tissue-specific Expression and Co-expression Analysis

The Jensen–Shannon (JS) divergence method (Cabili et al., 2011) was used for scoring tissue-specific expression. The *e*^t^ represented a standard expression mode that expressed only in one tissue, while the calculated transcript was defined as *e*. The distance between the two expression modes was calculated as follows. Tissue specificity score is defined as 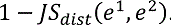.

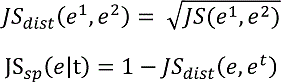

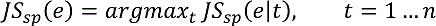

Co-expression analysis was analyzed by R package weighted correlation network analysis (WGCNA) (Zhang and Horvath, 2005). The lncRNAs with FPKM ≥1.5 and PCGs with FPKM ≥ 10 in any tissues were retained.

### TSS and TES Analysis

Based on the analysis of CAGE-seq and PolyA-seq, dominant TSS/TES with TPM ≥0.5 were used for further study. The TSS/TES frequency was defined as the average number of TSS/TES per locus. TSS/TES switching between any two tissues were considered as TSS/TES selection. For screening lncRNA TSS/TES motifs, known motifs including TATA, lnr, Y for TSS and PAS, U-rich for TES of PCGs and novel motifs were examined and identified by Homer and MEME, respectively. RNAfold was used to predict secondary structures of lncRNAs (Lorenz et al., 2016).

### Single Nucleotide Polymorphism (SNP) and GWAS Analysis

SNP data were collected from a previous study (Du et al., 2018). GWAS sites were collected from studies in *G. arboreum* (Du et al., 2018) and *G. hirsutum* (Li et al., 2018)(Wang et al., 2017)(Ma et al., 2018). GWAS sites in the At-subgenome of *G. hirsutum* were transformed into *G. arboreum*. Annovar (Wang et al., 2010) was used to detect the overlap between lncRNAs and SNP/GWAS sites. The LD values between SNPs were calculated by PLINK (1.9b) (Purcell et al., 2007). The haplotype of each *G. arboreum* line was identified by Beagle (5.1.24) (Browning and Browning, 2007). When investigating the seed phenotypes, we removed 17 lines with heterozygous genotype or without phenotypic data.

### ChIRP-seq

ChIRP-seq was performed according to previous reports (Percharde et al., 2018)(Zhao et al., 2018b) with some modification. Briefly, ovules (0 DPA) were crosslinked for 30 minutes in 0.4 M Sucrose, 10 mM Tris-HCl pH8.0, 5 mM β-ME, 1 % formaldehyde, protease inhibitor cocktail (Roche, Germany), RNase inhibitor, and then quenched with 0.125 M glycine for 5 min. The chromatin was sonicated into fragments of less than 300 bp. RNase treated chromatin was used as negative control. To capture the lncRNAs, 1 mL chromatin fragment was incubated with 100 pmol biotinylated lncRNA or LacZ probes in 2 mL solution (750 mM NaCl, 1% SDS, 50 mM Tris-HCl pH 7, 1 mM EDTA, 15% formamide, protease inhibitor cocktail, RNase inhibitor) at 37 ℃ overnight. Streptavidin-beads were added and incubated at room temperature for 30 minutes, and then were washed with 2× SSC, 0.5 % SDS and 1 mM PMSF for 5 times at 37 ℃. The precipitation was divided two parts: 10% and 90% were performed quantification assay for RNA and DNA enrichment, respectively. The percentage of retrieved RNA/DNA to input was used to reflect RNA/DNA enrichment by using qRT/q-PCR. The purified DNA fragments were used to build the sequencing library.

The clean data were mapped by STAR with EndToEnd. After removing PCR duplicates by picard, the surplus alignment results were delivered to MACS2 (Zhang et al., 2008) to call peaks. High-credibility peaks were filtered against its corresponding input with *p*-value cutoff 1e-5. The repeatability of ChIRP-seq was measured by calculating the Pearson’s correlation coefficient of the enrichment of common peaks identified in two replicates. By using CHIPseeker package (Yu et al., 2015) in R, we offered every peak an annotation with the definition for a promoter being 3 Kb around the TSS. GO enrichment analysis for PCGs with high-credibility peak was finished by ChIP-Enrich (Welch et al., 2014) with *p*-value cutoff 0.05. Motif analysis was executed with sequences of all peaks within +/-50 bp around peak summits. Only motifs with high significance were shown.

### VIGS Assay

VIGS assay was performed as previously reported (Gao et al., 2013). The 699^th^-1295^th^ bp of *lnc-Ga13g0352* which has lower sequence similarity than other regions of the *Gossypium arboreum* transcriptome was amplified and ligated to pTRV2 vectors, and then introduced into *Agrobacterium tumefaciens* strain GV3101. The transformed *Agrobacterium* strains were resuspended with infiltration buffer (10 mM MgCl_2_, 10 mM MES and 200 mΜ acetosyringone). The transformant with pTRV2: *lnc-Ga13g0352* or pTRV2: 00 (empty vector) were mixed with pTRV1 in a 1:1 ratio and infiltrated into cotyledons of seedlings (two weeks). The injected seedlings continued to grow in growth chamber at 25 ℃ for another two weeks. For detecting the effect of VIGS, the true leaves were harvested for total RNA isolation. The oligo dT was used as the reverse transcription primer of cDNA synthesis for qRT-PCR.

## Accession numbers

Sequence data can be found in NCBI Sequence Read Archive under the accession numbers PRJNA542206, PRJNA507565and PRJNA373801 (details provided in Supplemental Table S9).

## SUPPLEMENTAL DATA

The following supplemental materials are available.

Supplemental Table S1. Genomic coordinates of lncRNAs in BED12 format and category of lncRNAs based on terminal signal, genome location and miRNA in Gossypium arboreum.

Supplemental Table S2. The expression (FPKM) of lncRNAs across 21 tissues.

Supplemental Table S3. The expression (FPKM) of PCGs across 21 tissues.

Supplemental Table S4. The corresponding number IDs of tissues.

Supplemental Table S5. The genomic locations of lncRNAs which were related TSS switches of PCGs.

Supplemental Table S6. The WGCNA analysis on the expression of lncRNAs and PCGs.

Supplemental Table S7. The ChIRP-seq peaks and associated gene information.

Supplemental Table S8. All primers used in this study.

Supplemental Table S9. Sources and information of NGS datasets used in this study.

Supplemental Table S10. Information of software and programs used in this study.

## ACKNOWLEDGEMENTS

We thank Prof. Xueying Guan from Zhejiang University (Hangzhou, China) for providing the VIGS vectors.

---

**Supplemental Figure S1.**
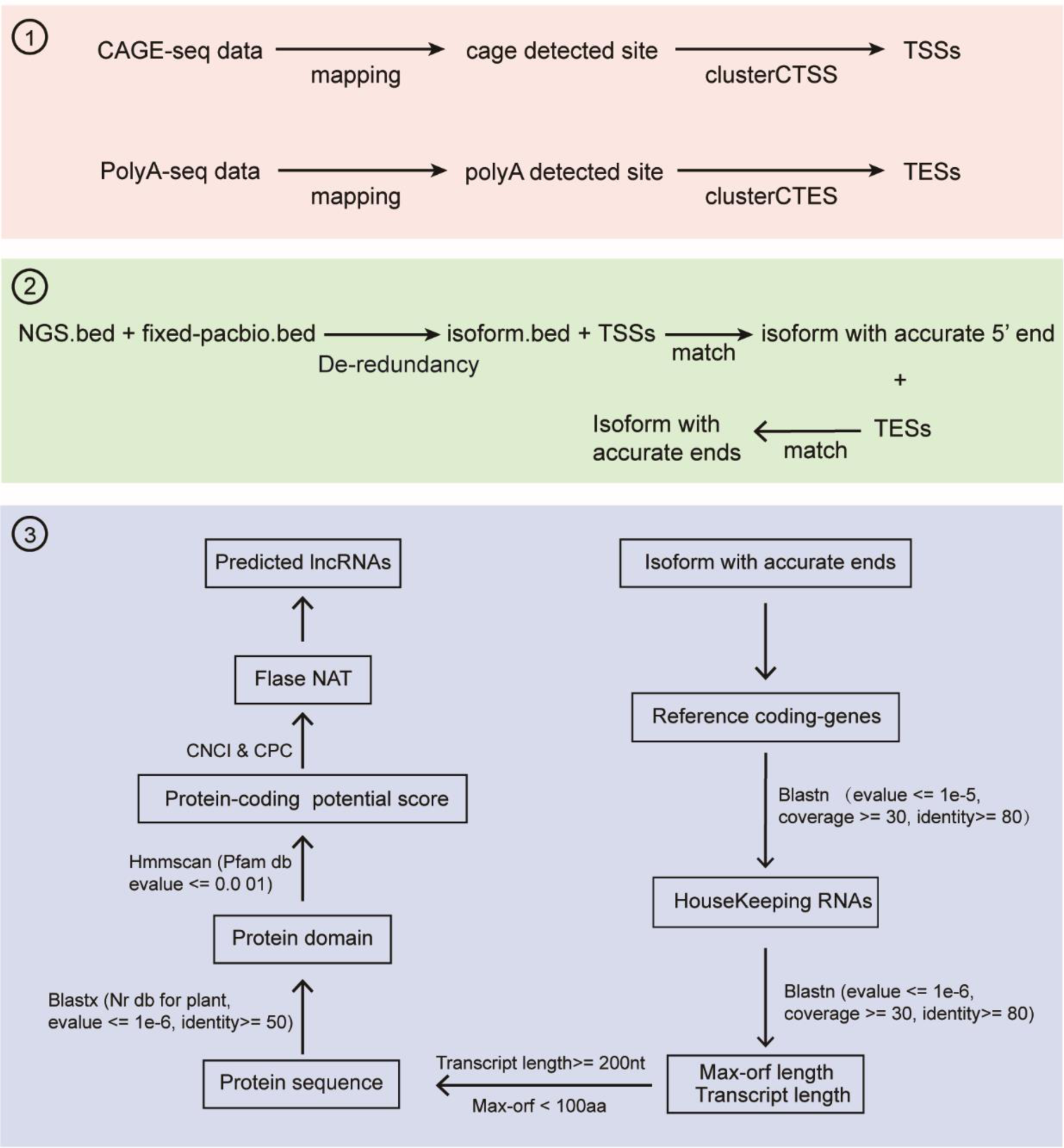
The detailed information of three steps of PULL pipeline. Steps 1-3 omitted in Figure 1b were further described. Step 1 represents processing of terminal sequencing data including CAGE-Seq and PolyA-seq. Step 2 represents processing of transcript sequencing data including ssRNA-Seq and Iso-seq. Step 3 represents processing of lncRNA identification.

**Supplemental Figure S2.**
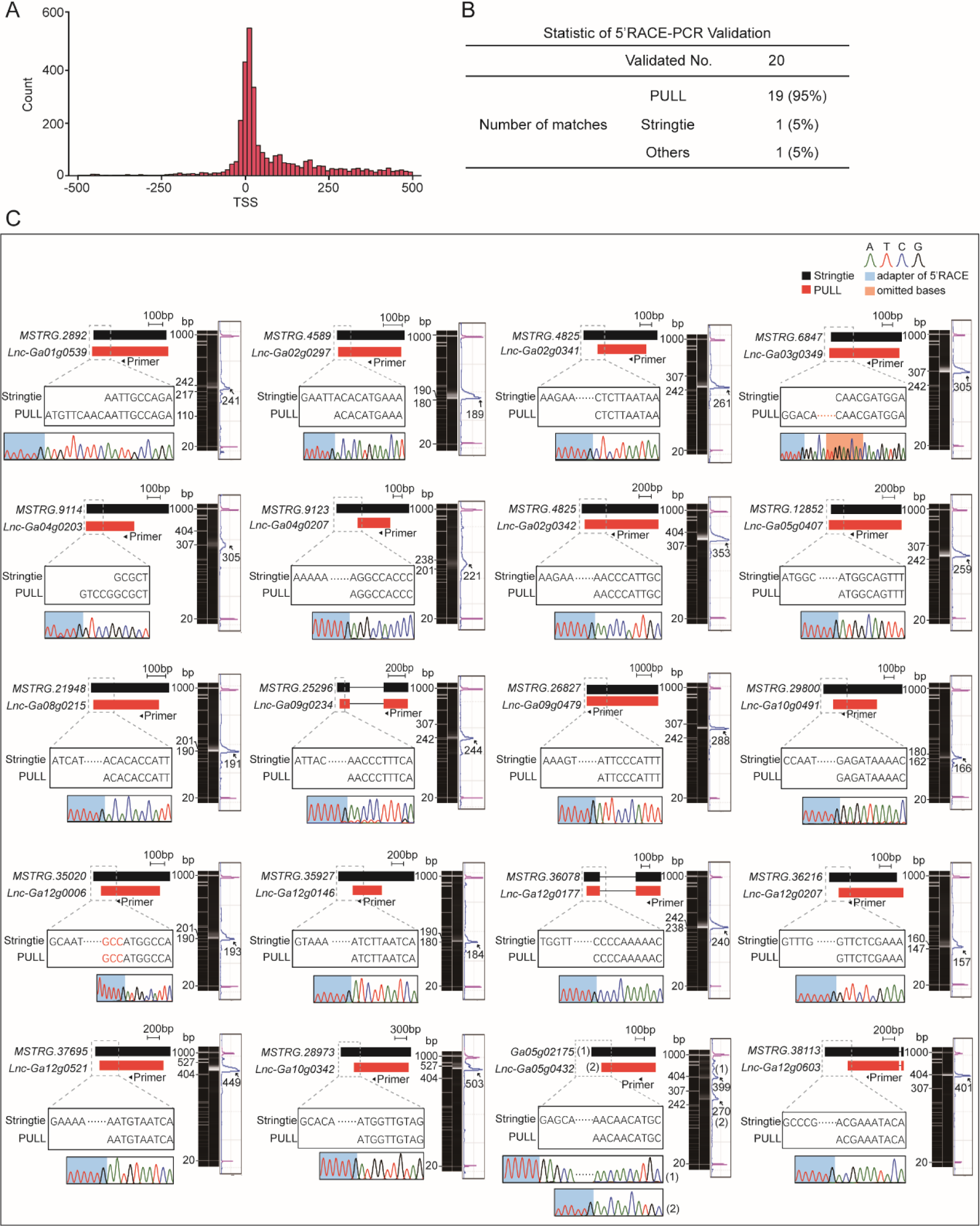
The Sanger sequencing validation of 5’ ends signals of lncRNAs predicted by PULL. A, The distribution of distance between 5’ ends identified by StringTie and PULL. The ordinate represents the number of transcripts, and the abscissa represents the distance. B, The statistic of 5’ RACE experiment for lncRNAs. C, The Sanger sequencing results of 5’ RACE experiment for 20 lncRNAs.

**Supplemental Figure S3.**
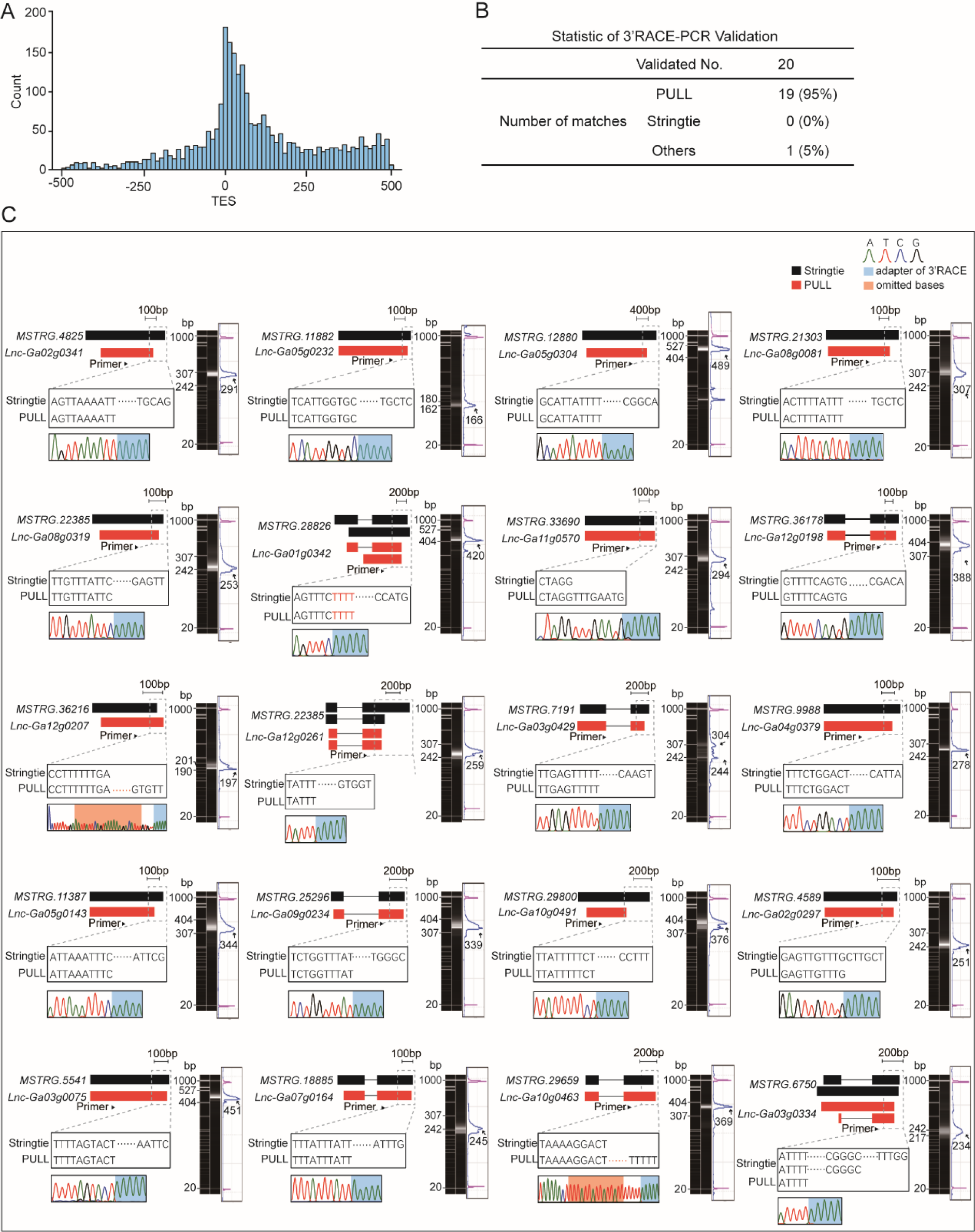
The Sanger sequencing validation of 3’ ends signals of lncRNAs predicted by PULL. A, The distribution of distance between 3’ ends identified by StringTie and PULL. The ordinate represents the number of transcripts, and the abscissa represents the distance. B, The statistic of 3’ RACE experiment for lncRNAs. C, The Sanger sequencing results of 3’ RACE experiment for 20 lncRNAs.

**Supplemental Figure S4.**
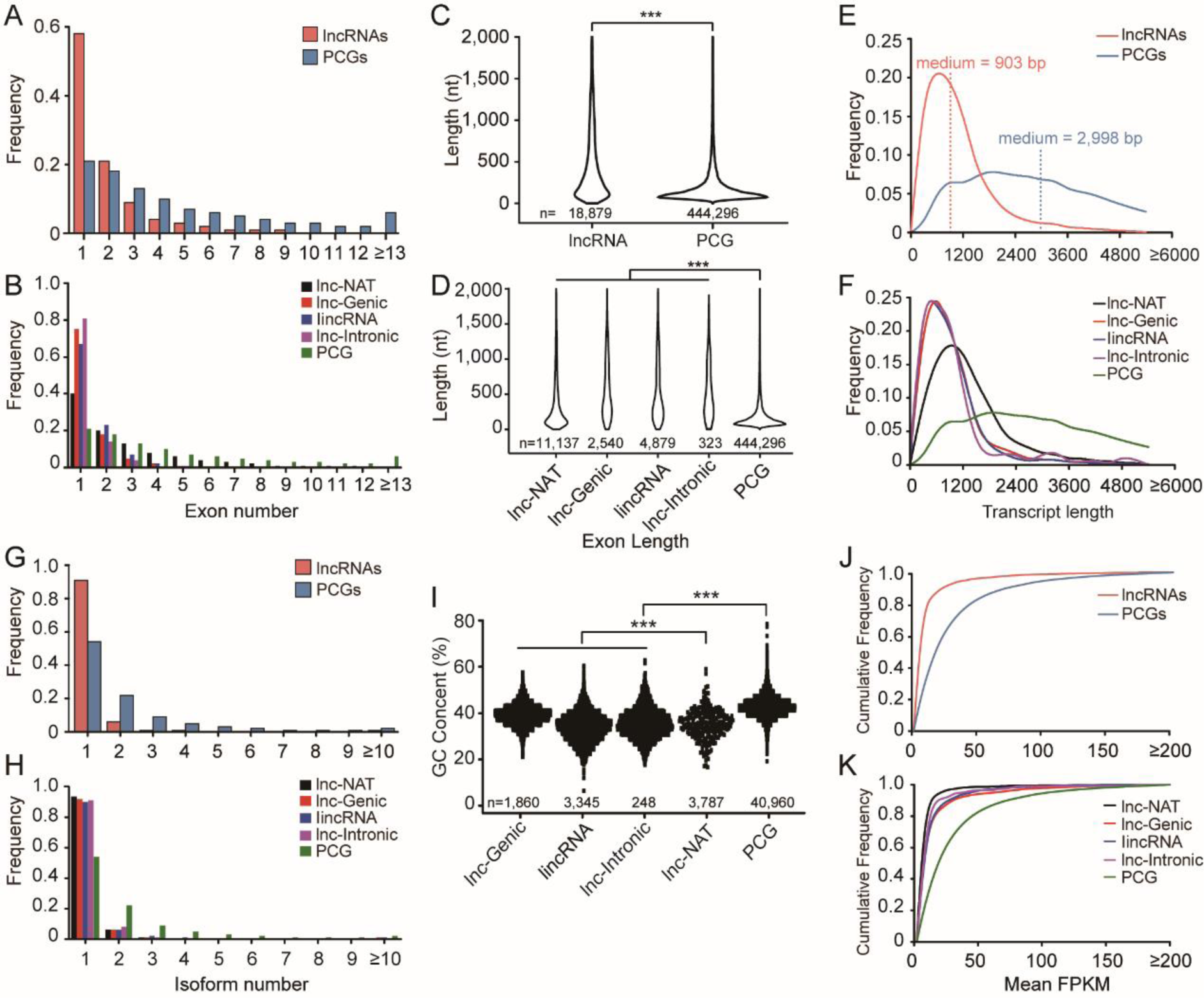
The genomic features of lncRNAs and PCGs in G. arboreum. The comparison of exon number A-B, exon length C-D, transcript length E-F, isoform number G-H, GC content I, and cumulative expression J-K, for lncRNAs and PCGs. The Mann-Whitney test were used for significant differences (*** P value < 0.01).

**Supplemental Figure S5.**
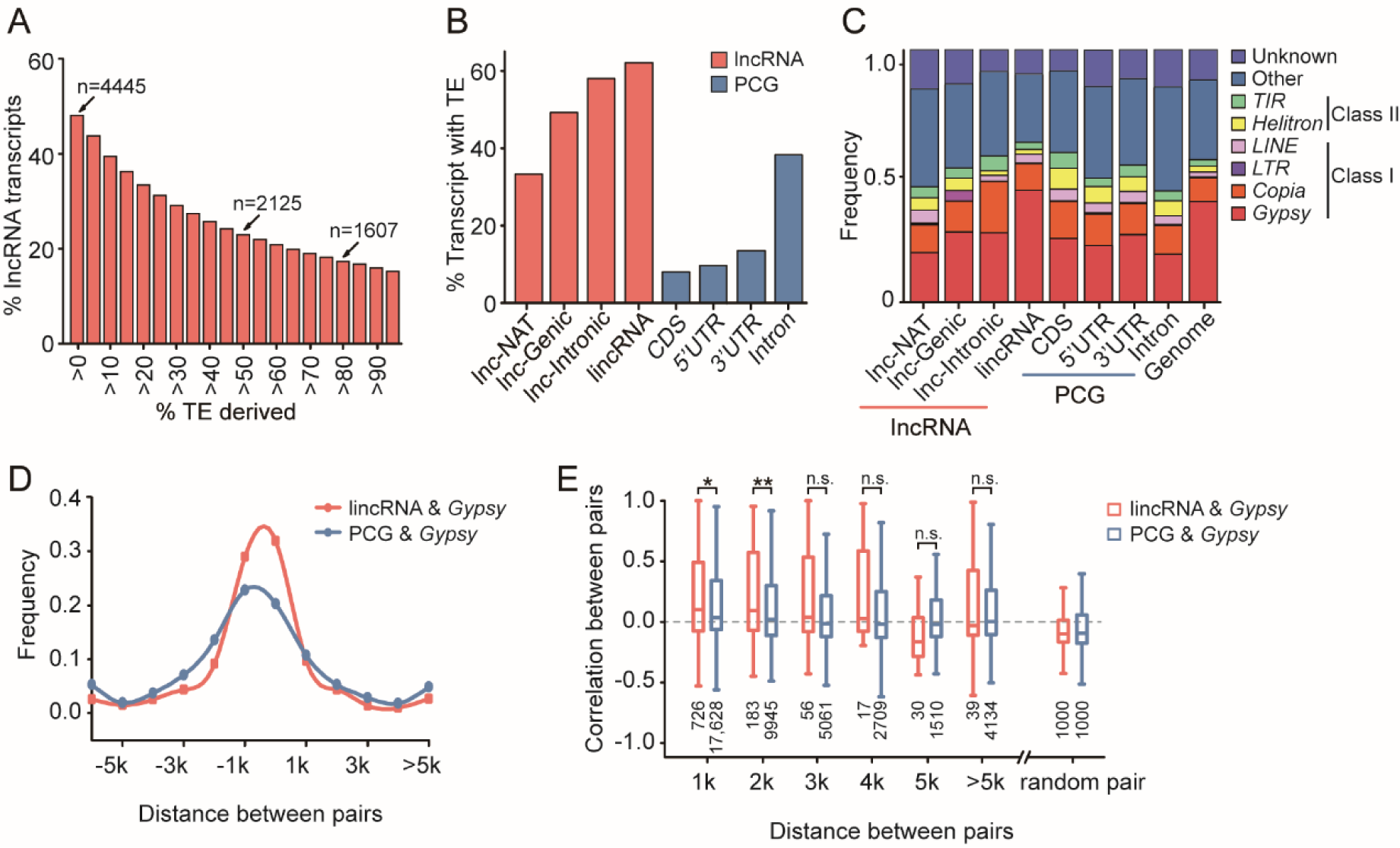
TEs might contribute to the sequence composition and expressional regulation of lncRNA in G. arboreum. A, Distribution of lncRNA transcripts derived from TEs (> 0 % to > 95 %). The number of transcripts with > 0 %, > 50 % and > 80 % TE-derived DNA exons were indicated above the bars. B, Percentage of lncRNA transcripts with at least one exon overlapping with TEs (more than 10 bp). C, TE components in lncRNAs, PCGs, and the whole genome. D, Distribution frequency of the distances from lincRNA and PCG to their closest Gypsy. E, Expression correlation between lincRNA, PCG, and their closest Gypsy. The random pair represents the expression correlation for random pairing between lncRNA or PCG and Gypsy. The significance levels are indicated by asterisks (Mann-Whitney test, * P value < 0.05; ** P value < 0.01; n.s. represents not significant). The number of sample size (n) for analysis are indicated below the boxplot.

**Supplemental Figure S6.**
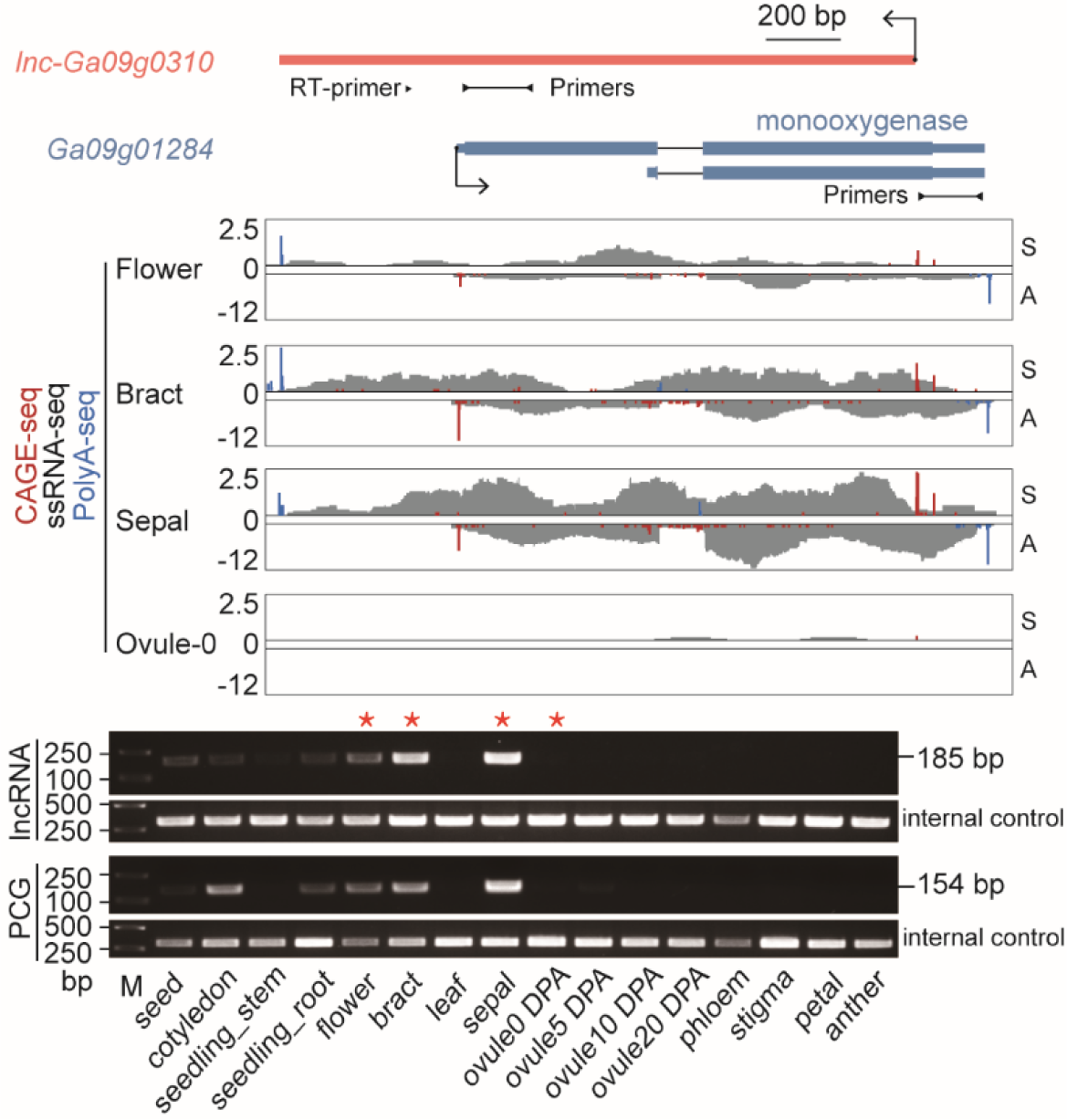
The co-expression of a lnc-NAT with its cognate antisense PCG. Top: The genomic locations of the lnc-NAT and PCG. The primers for reverse transcription (RT-primer) and qPCR primer pairs (Primers) are indicated. Middle: CAGE-seq, PolyA-seq and ssRNA-seq signals in four tissues. S and A represent the sense and antisense strands, respectively. Bottom: validation of semi-RT-PCR across tissues. Red stars indicate the four tissues with RNA-seq data shown above.

**Supplemental Figure S7.**
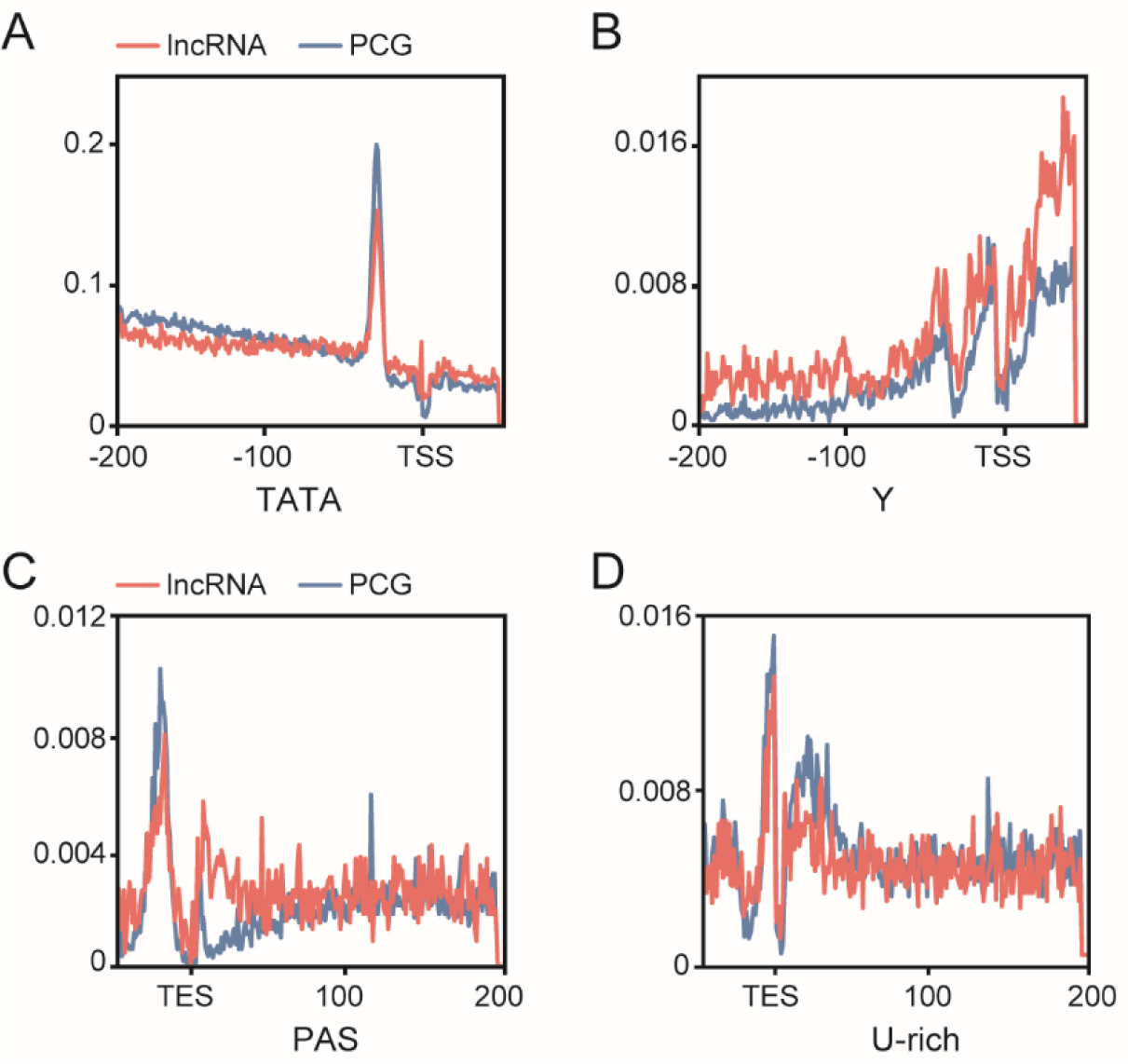
The canonical cis elements of Pol II transcription found in lncRNAs and PCGs. A-B, The distribution of TATA and Y around the TSSs of lncRNAs and PCGs. C-D, The distribution of PAS and U-rich around the TESs of lncRNAs and PCGs. The abscissa represents the distance from TSS/TES, and the ordinate represents frequency of the element at the corresponding site.

**Supplemental Figure S8.**
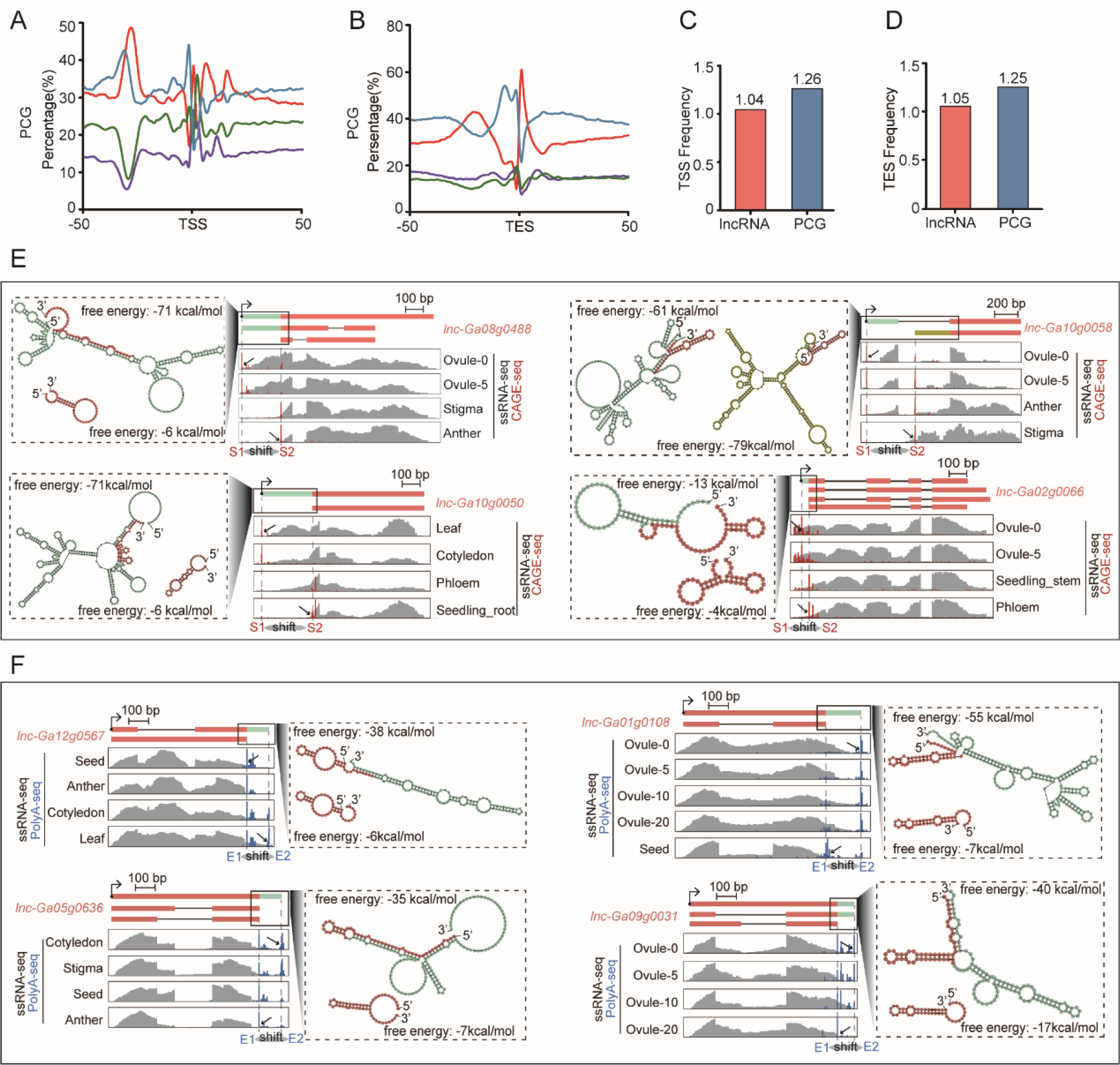
The switches of TSS and TES and their effect on the secondary structure of lncRNAs. A, The nucleotide composition around the TSS of lncRNAs and PCGs. B, The nucleotide composition around the TES of lncRNAs and PCGs. C, TSS frequency of lncRNAs and PCGs. D, TES frequency of lncRNAs and PCGs. E, The changes of RNA secondary structure caused by TSS switches. F, The changes of RNA secondary structure caused by alternative TES switches.

**Supplemental Figure S9.**
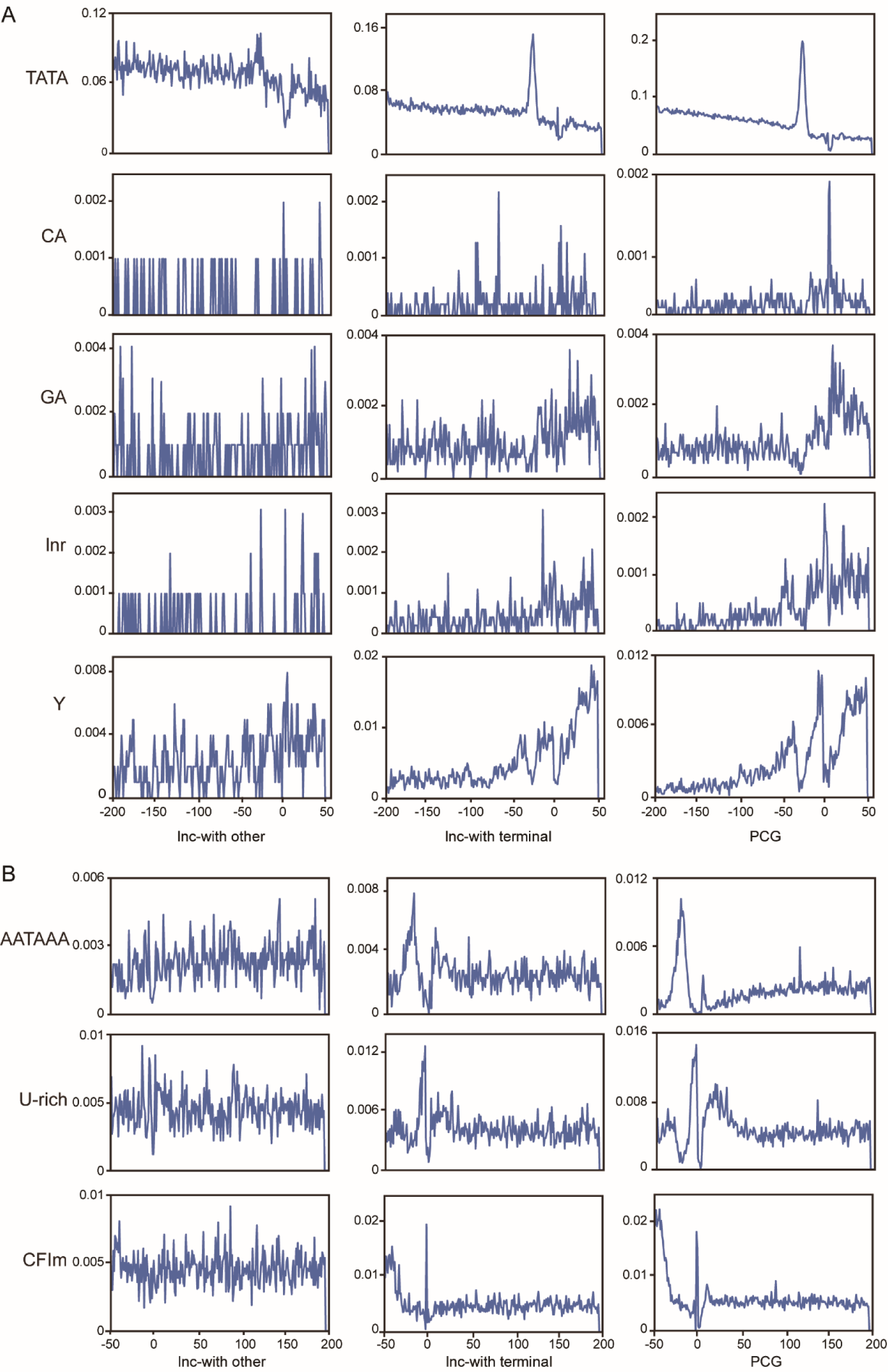
The distribution of known motifs in promoter and terminator of Pol II in different types of lncRNAs. A, The known cis-elements in the promoter regions. B, The known cis-elements in the terminator regions.

**Supplemental Figure S10.**
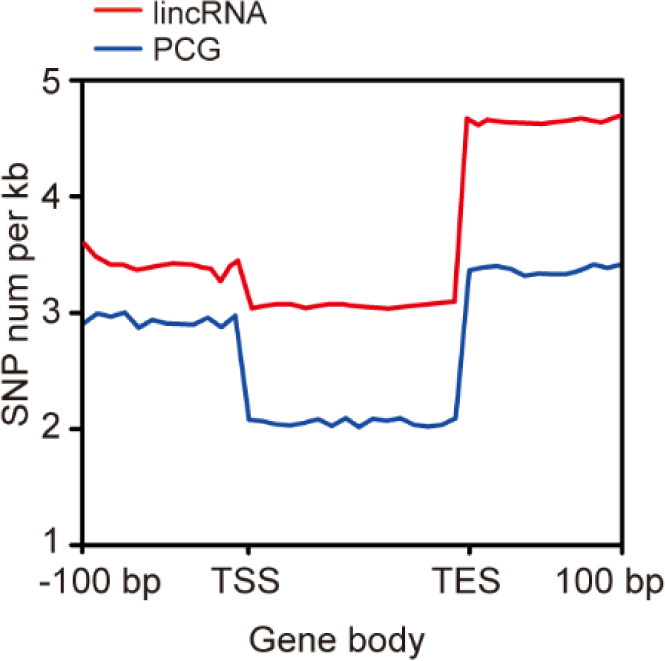
SNP frequency distribution on the gene body of lincRNAs (red line) and PCGs (blue line). 100 bp upstream of TSS, gene body and 100 bp downstream of TES were shown as abscissa.

**Supplemental Figure S11.**
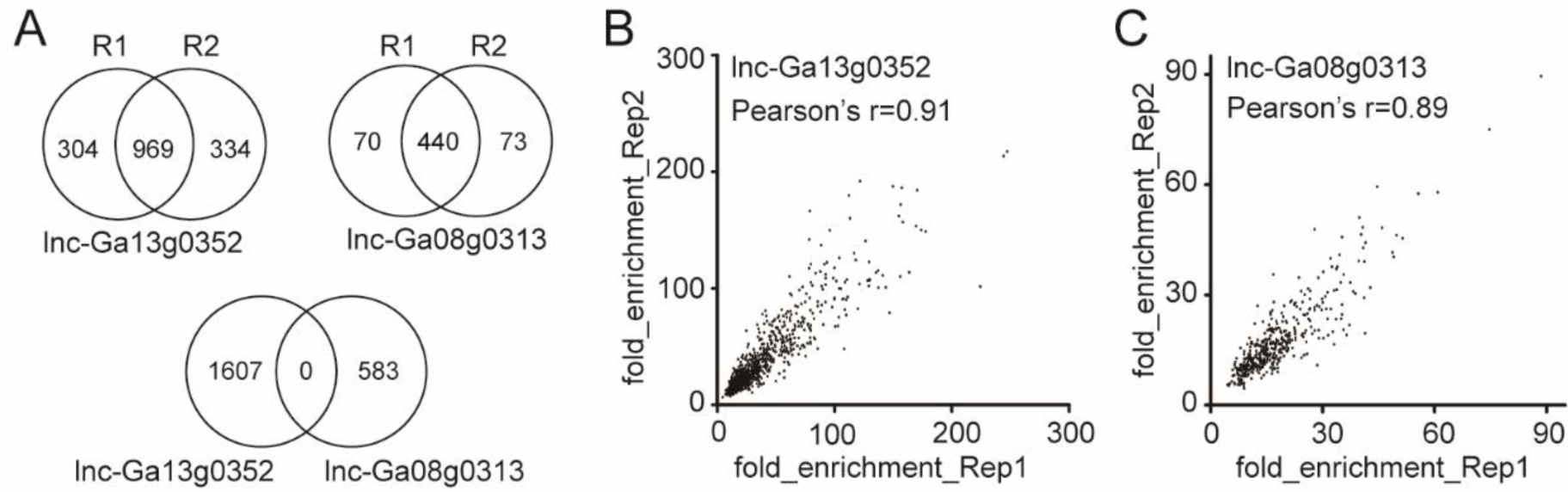
The evaluation on the biological repeatability of ChIRP-seq for lnc-Ga13g0352 and lnc-Ga08g0313. A, The number of binding peaks of lnc-Ga13g0352 and lnc-Ga08g0313 in two biological replicates (Top). There is no any overlapping peak between two lncRNAs’ binding sites (Bottom). R1 and R2 represent the two biological replicates. B-C, The plots of peak-enriching fold between two replicates for lnc-Ga13g0352 (B) and lnc-Ga08g0313 (C), in which the pearson’s correlation coefficient are shown.

**Supplemental Figure S12.**
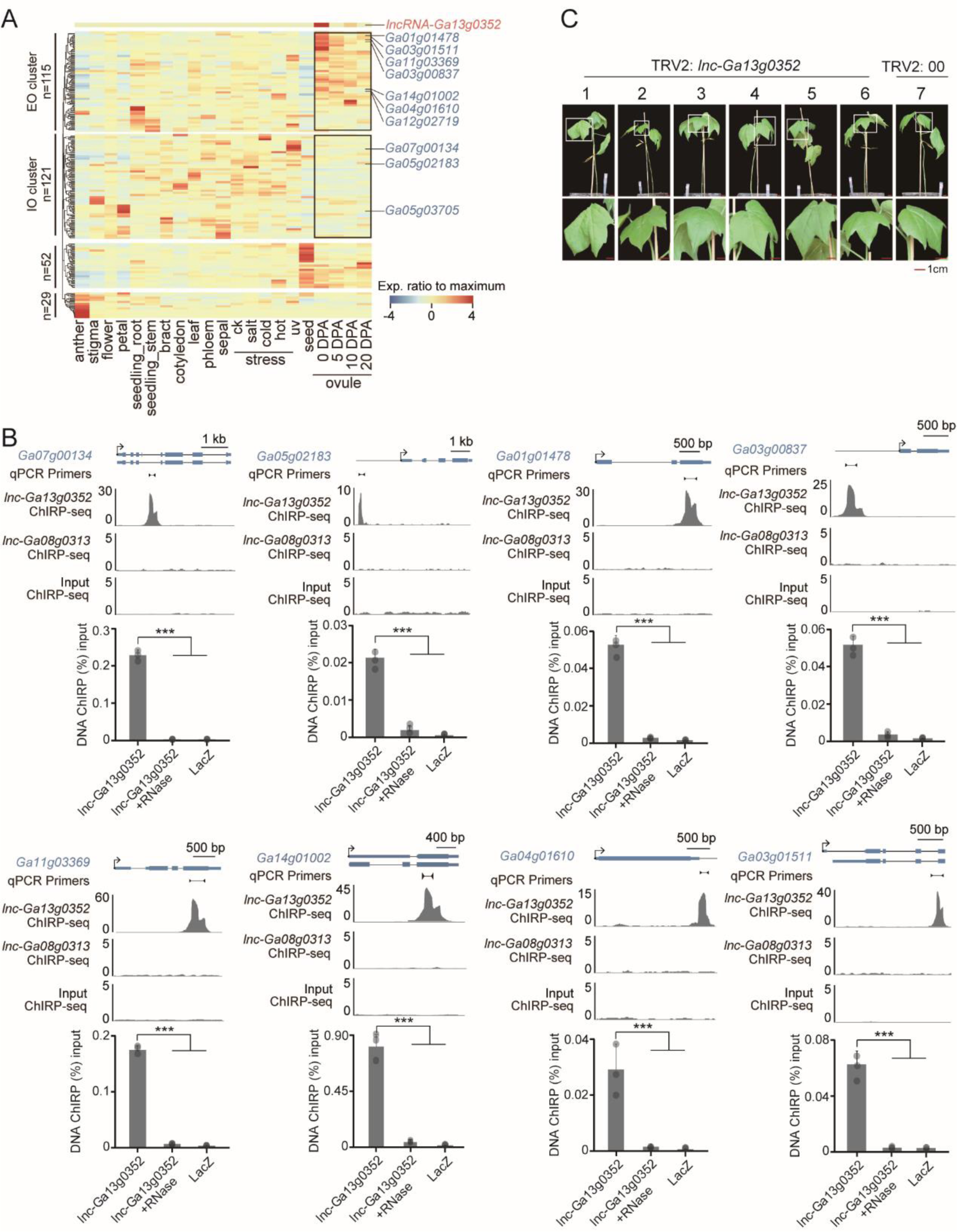
The validation of lnc-Ga13g0352 targeting PCGs and phenotypic observation in VIGS. A, The expression and clustering of the target PCGs of lnc-Ga13g0352. The common name of some homologs for known PCGs were indicated. B, The ChIRP-seq peaks and validation with ChIRP-qPCR for the targeting genes of lnc-Ga13g0352. The peaks from one biological replicate were shown. The lacZ and lnc-Ga08g0313 probes were used as non-targeting control, and RNase-treated samples were used as negative control for ChIRP. Black line with triangles represent pairs of qPCR primers. The significance are indicated by asterisks (two-tailed t-test, three biological replicates, error bar represents standard deviation, *** P value <0.001). C, The phenotypic observation of cotton seedlings in VIGS assay (two weeks after infiltration), indicating there is no visible effect on the normal growth.

**Supplemental Figure S13.**
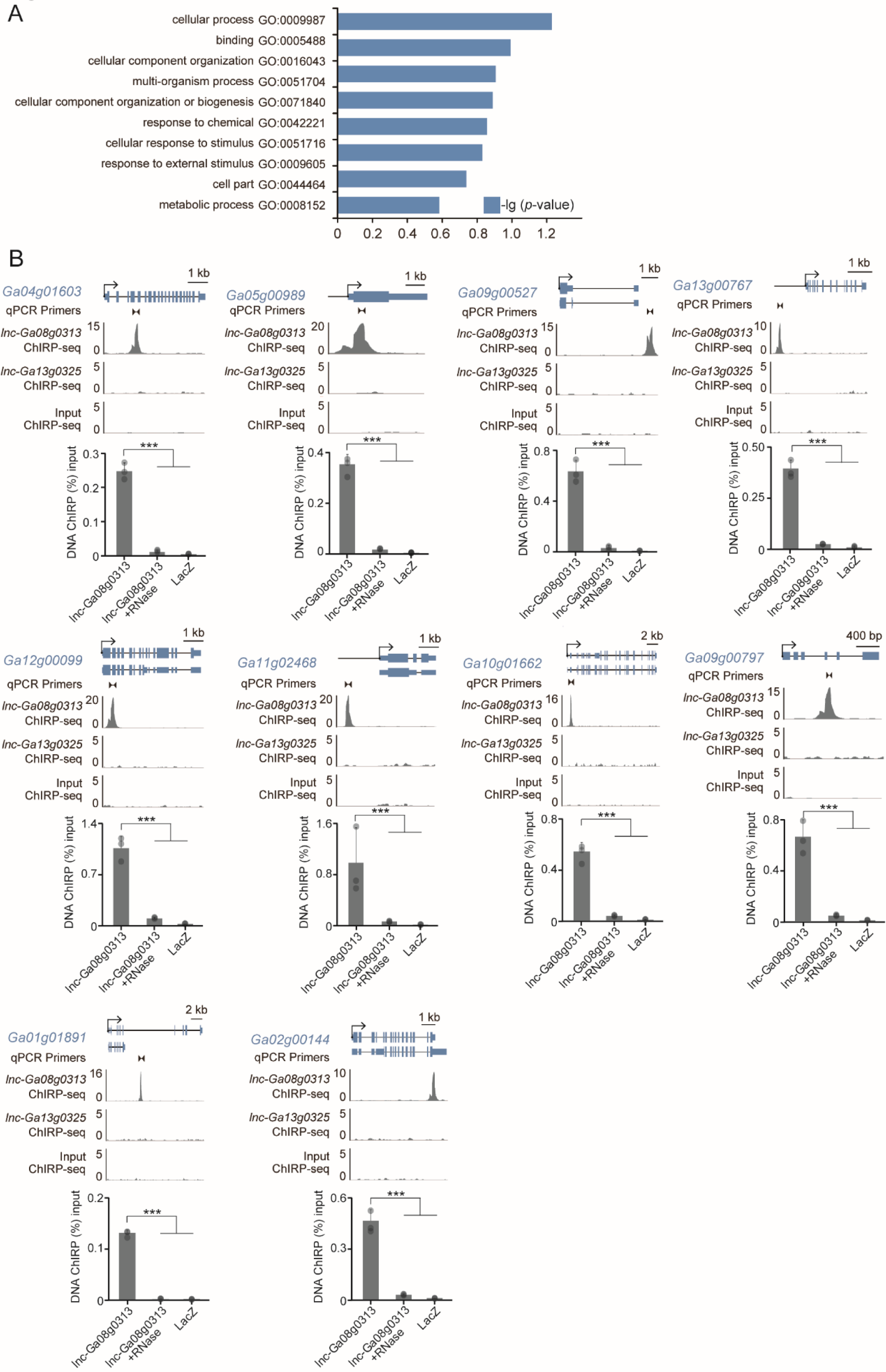
The analysis and validation of lnc-Ga08g0313 targeting genes. A, Top 10 GO terms in GO enrichment analysis for lnc-Ga08g0313 binding genes. B, The ChIRP-seq peaks and ChIRP-qPCR for the targeting genes of lncRNAs. The peaks from one biological replicate were shown. The lacZ and lnc-Ga13g0352 probes were used as non-targeting control, and RNase-treated samples were used as negative control for ChIRP. Black line with triangles represent pairs of qPCR primers. The significance levels are indicated by asterisks (two-tailed t-test, three biological replications, error bars represent standard deviation, *** P value < 0.001).

## References

1. Ariel F, Jegu T, Latrasse D, Romero-Barrios N, Christ A, Benhamed M, Crespi M (2014) Noncoding transcription by alternative RNA polymerases dynamically regulates an auxin-driven chromatin loop. Mol Cell 55: 383–96

2. Bardou F, Ariel F, Simpson CG, Romero-Barrios N, Laporte P, Balzergue S, Brown JWS, Crespi M (2014) Long Noncoding RNA Modulates Alternative Splicing Regulators in Arabidopsis. Dev Cell 30: 166–176

3. Boley N, Stoiber MH, Booth BW, Wan KH, Hoskins RA, Bickel PJ, Celniker SE, Brown JB (2014) Genome-guided transcript assembly by integrative analysis of RNA sequence data. Nat Biotechnol 32: 341–346

4. Browning SR, Browning BL (2007) Rapid and accurate haplotype phasing and missing-data inference for whole-genome association studies by use of localized haplotype clustering. Am J Hum Genet 81: 1084–1097

5. Cabili MN, Trapnell C, Goff L, Koziol M, Tazon-Vega B, Regev A, Rinn JL (2011) Integrative annotation of human large intergenic noncoding RNAs reveals global properties and specific subclasses. Genes Dev 25: 1915–27

6. Csorba T, Questa JI, Sun Q, Dean C (2014) Antisense COOLAIR mediates the coordinated switching of chromatin states at FLC during vernalization. Proc Natl Acad Sci U S A 111: 16160–5

7. Dai X, Zhuang Z, Zhao PX (2018) PsRNATarget: Aplant small RNAtarget analysis server (2017 release). Nucleic Acids Res 46: W49–W54

8. Deng P, Liu S, Nie X, Weining S, Wu L (2018) Conservation analysis of long non-coding RNAs in plants. Sci China Life Sci 61: 190–198

9. Ding J, Lu Q, Ouyang Y, Mao H, Zhang P, Yao J, Xu C, Li X, Xiao J, Zhang Q (2012) Along noncoding RNAregulates photoperiod-sensitive male sterility, an essential component of hybrid rice. Proc Natl Acad Sci U S A 109: 2654–9

10. Dobin A, Davis CA, Schlesinger F, Drenkow J, Zaleski C, Jha S, Batut P, Chaisson M, Gingeras TR (2013) STAR: Ultrafast universal RNA-seq aligner. Bioinformatics 29: 15–21

11. Du X, Huang G, He S, Yang Z, Sun G, Ma X, Li N, Zhang X, Sun J, Liu M, et al (2018) Resequencing of 243 diploid cotton accessions based on an updated Agenome identifies the genetic basis of key agronomic traits. Nat Genet 50: 796–802

12. Engreitz JM, Haines JE, Perez EM, Munson G, Chen J, Kane M, Mcdonel PE, Guttman M, Lander ES (2016) Local regulation of gene expression by lncRNApromoters, transcription and splicing. Nature 539: 452–455

13. Fang L, Wang Q, Hu Y, Jia Y, Chen J, Liu B, Zhang Z, Guan X, Chen S, Zhou B, et al (2017) Genomic analyses in cotton identify signatures of selection and loci associated with fiber quality and yield traits. Nat Genet. doi: 10.1038/ng.3887

14. Franco-Zorrilla JM, Valli A, Todesco M, Mateos I, Puga MI, Rubio-Somoza I, Leyva A, Weigel D, García JA, Paz-Ares J (2007) Target mimicry provides a new mechanism for regulation of microRNA activity. Nat Genet 39: 1033–1037

15. Fu H, Yang D, Su W, Ma L, Shen Y, Ji G, Ye X, Wu X, Li QQ (2016) Genome-wide dynamics of alternative polyadenylation in rice. Genome Res 26: 1753–1760

16. Gao X, Li F, Li M, Kianinejad AS, Dever JK, Wheeler TA, Li Z, He P, Shan L (2013) Cotton GhBAK1 Mediates Verticillium Wilt Resistance and Cell Death. J Integr Plant Biol 55: 586–596

17. Golicz AA, Singh MB, Bhalla PL (2018) The Long Intergenic Noncoding RNA(LincRNA) Landscape of the Soybean Genome. Plant Physiol 176: 2133–2147

18. Haberle V, Forrest ARR, Hayashizaki Y, Carninci P, Lenhard B (2015) CAGEr: precise TSS data retrieval and high-resolution promoterome mining for integrative analyses. Nucleic Acids Res 43: e51–e51

19. Heinz S, Benner C, Spann N, Bertolino E, Lin YC, Laslo P, Cheng JX, Murre C, Singh H, Glass CK (2010) Simple Combinations of Lineage-Determining Transcription Factors Prime cis-Regulatory Elements Required for Macrophage and B Cell Identities. Mol Cell 38: 576–589

20. Hon C-C, Ramilowski JA, Harshbarger J, Bertin N, Rackham OJL, Gough J, Denisenko E, Schmeier S, Poulsen TM, Severin J, et al (2017) An atlas of human long non-coding RNAs with accurate 5′ ends. Nature 543: 199–204

21. Hou S, Zhu G, Li Y, Li W, Fu J, Niu E, Li L, Zhang D, Guo W(2018) Genome-Wide Association Studies Reveal Genetic Variation and Candidate Genes of Drought Stress Related Traits in Cotton (Gossypium hirsutum L.). Front Plant Sci 9: 1276

22. Kalvari I, Argasinska J, Quinones-Olvera N, Nawrocki EP, Rivas E, Eddy SR, Bateman A, Finn RD, Petrov AI (2018) Rfam 13.0: shifting to a genome-centric resource for non-coding RNAfamilies. Nucleic Acids Res 46: D335–D342

23. Kang Y-J, Yang D-C, Kong L, Hou M, Meng Y-Q, Wei L, Gao G (2017) CPC2: a fast and accurate coding potential calculator based on sequence intrinsic features. Nucleic Acids Res 45: W12–W16

24. Kawaji H, Lizio M, Itoh M, Kanamori-Katayama M, Kaiho A, Nishiyori-Sueki H, Shin JW, Kojima-Ishiyama M, Kawano M, Murata M, et al (2014) Comparison of CAGE and RNA-seq transcriptome profiling using clonally amplified and single-molecule next-generation sequencing. Genome Res 24: 708–17

25. Kindgren P, Ard R, Ivanov M, Marquardt S (2018) Transcriptional read-through of the long non-coding RNASVALKA governs plant cold acclimation. Nat Commun 9: 4561

26. Li C, Fu Y, Sun R, Wang Y, Wang Q (2018) Single-Locus and Multi-Locus Genome-Wide Association Studies in the Genetic Dissection of Fiber Quality Traits in Upland Cotton (Gossypium hirsutum L.). Front Plant Sci 9: 1083

27. Li F, Fan G, Wang K, Sun F, Yuan Y, Song G, Li Q, Ma Z, Lu C, Zou C, et al (2014a) Genome sequence of the cultivated cotton Gossypium arboreum. Nat Genet 46: 567–72

28. Li L, Eichten SR, Shimizu R, Petsch K, Yeh C-T, Wu W, Chettoor AM, Givan SA, Cole RA, Fowler JE, et al (2014b) Genome-wide discovery and characterization of maize long non-coding RNAs. Genome Biol 15: R40

29. Lisch D (2013) How important are transposons for plant evolution? Nat Rev Genet 14: 49–61

30. Liu J, Jung C, Xu J, Wang H, Deng S, Bernad L, Arenas-Huertero C, Chua N-H (2012) Genome-wide analysis uncovers regulation of long intergenic noncoding RNAs in Arabidopsis. Plant Cell 24: 4333–45

31. Liu X, Hao L, Li D, Zhu L, Hu S (2015) Long Non-coding RNAs and Their Biological Roles in Plants. Genomics Proteomics Bioinformatics 13: 137–147

32. Lorenz R, Bernhart SH, Höner zu Siederdissen C, Tafer H, Flamm C, Stadler PF, Hofacker IL (2011) ViennaRNAPackage 2.0. Algorithms Mol Biol 6: 26

33. Lorenz R, Hofacker IL, Stadler PF (2016) RNAfolding with hard and soft constraints. Algorithms Mol Biol 11: 8

34. Ma Z, He S, Wang X, Sun J, Zhang Y, Zhang G, Wu L, Li Z, Liu Z, Sun G, et al (2018) Resequencing a core collection of upland cotton identifies genomic variation and loci influencing fiber quality and yield. Nat Genet 50: 803–813

35. Mercer TR, Dinger ME, Mattick JS (2009) Long non-coding RNAs: insights into functions. Nat Rev Genet 10: 155–159

36. Paytuví Gallart A, Hermoso Pulido A, Anzar Martínez de Lagrán I, Sanseverino W, Aiese Cigliano R (2016) GREENC: a Wiki-based database of plant lncRNAs. Nucleic Acids Res 44: D1161-6

37. Percharde M, Lin CJ, Yin Y, Guan J, Peixoto GA, Bulut-Karslioglu A, Biechele S, Huang B, Shen X, Ramalho-Santos M (2018) ALINE1-Nucleolin Partnership Regulates Early Development and ESC Identity. Cell 174: 391-405.e19

38. Pertea M, Pertea GM, Antonescu CM, Chang T-C, Mendell JT, Salzberg SL (2015) StringTie enables improved reconstruction of a transcriptome from RNA-seq reads. Nat Biotechnol 33: 290–295

39. Purcell S, Neale B, Todd-Brown K, Thomas L, Ferreira MAR, Bender D, Maller J, Sklar P, De Bakker PIW, Daly MJ, et al (2007) PLINK: A tool set for whole-genome association and population-based linkage analyses. Am J Hum Genet 81: 559–575

40. Quinlan AR, Hall IM (2010) BEDTools: a flexible suite of utilities for comparing genomic features. Bioinformatics 26: 841–842

41. Ransohoff JD, Wei Y, Khavari PA(2017) The functions and unique features of long intergenic non-coding RNA. Nat Rev Mol Cell Biol 19: 143–157

42. Shan C-M, Shangguan X-X, Zhao B, Zhang X-F, Chao L, Yang C-Q, Wang L-J, Zhu H-Y, Zeng Y-D, Guo W-Z, et al (2014) Control of cotton fibre elongation by a homeodomain transcription factor GhHOX3. Nat Commun 5: 5519

43. Sigman MJ, Slotkin RK (2016) The first rule of plant transposable element silencing: Location, location, location. Plant Cell 28: 304–313

44. StLaurent G, Wahlestedt C, Kapranov P (2015) The Landscape of long noncoding RNAclassification. Trends Genet 31: 249–251

45. Sun L, Luo H, Bu D, Zhao G, Yu K, Zhang C, Liu Y, Chen R, Zhao Y (2013) Utilizing sequence intrinsic composition to classify protein-coding and long non-coding transcripts. Nucleic Acids Res 41: e166–e166

46. Tokizawa M, Kusunoki K, Koyama H, Kurotani A, Sakurai T, Suzuki Y, Sakamoto T, Kurata T, Yamamoto YY (2017) Identification of Arabidopsis genic and non-genic promoters by paired-end sequencing of TSS tags. Plant J 90: 587–605

47. Uszczynska-Ratajczak B, Lagarde J, Frankish A, Guigó R, Johnson R (2018) Towards a complete map of the human long non-coding RNAtranscriptome. Nat Rev Genet 19: 535–548

48. Wang K, Huang G, Zhu Y (2016) Transposable elements play an important role during cotton genome evolution and fiber cell development. Sci China Life Sci 59: 112–121

49. Wang K, Li M, Hakonarson H (2010) ANNOVAR: functional annotation of genetic variants fromhigh-throughput sequencing data. Nucleic Acids Res 38: e164–e164

50. Wang K, Wang D, Zheng X, Qin A, Zhou J, Guo B, Chen Y, Wen X, Ye W, Zhou Y, et al (2019a) Multi-strategic RNA-seq analysis reveals a high-resolution transcriptional landscape in cotton. Nat Commun. doi: 10.1038/s41467-019-12575-x

51. Wang K, Wang D, Zheng X, Qin A, Zhou J, Guo B, Chen Y, Wen X, Ye W, Zhou Y, et al (2019b) Multi-strategic RNA-seq analysis reveals a high-resolution transcriptional landscape in cotton. Nat Commun 10: 4714

52. Wang M, Tu L, Lin M, Lin Z, Wang P, Yang Q, Ye Z, Shen C, Li J, Zhang L, et al (2017) Asymmetric subgenome selection and cis-regulatory divergence during cotton domestication. Nat Genet 49: 579–587

53. Wang M, Yuan D, Tu L, Gao W, He Y, Hu H, Wang P, Liu N, Lindsey K, Zhang X (2015a) Long noncoding RNAs and their proposed functions in fibre development of cotton (Gossypium spp.). New Phytol 207: 1181–1197

54. Wang M, Yuan D, Tu L, Gao W, He Y, Hu H, Wang P, Liu N, Lindsey K, Zhang X (2015b) Long noncoding RNAs and their proposed functions in fibre development of cotton (Gossypium spp.). New Phytol 207: 1181–1197

55. Wang R, Zheng D, Yehia G, Tian B (2018a) Acompendium of conserved cleavage and polyadenylation events in mammalian genes. Genome Res 28: 1427–1441

56. Wang Y, Luo X, Sun F, Hu J, Zha X, Su W, Yang J (2018b) Overexpressing lncRNALAIR increases grain yield and regulates neighbouring gene cluster expression in rice. Nat Commun 9: 3516

57. Welch RP, Lee C, Imbriano PM, Patil S, Weymouth TE, Smith RA, Scott LJ, Sartor MA(2014) ChIP-Enrich: gene set enrichment testing for ChIP-seq data. Nucleic Acids Res 42: e105–e105

58. Wu H, Yang L, Chen L-L (2017) The Diversity of Long Noncoding RNAs and Their Generation. Trends Genet 33: 540–552

59. Xu H, Luo X, Qian J, Pang X, Song J, Qian G, Chen J, Chen S (2012) FastUniq: AFast De Novo Duplicates Removal Tool for Paired Short Reads. PLoS One 7: e52249

60. Yamamoto YY, Yoshitsugu T, Sakurai T, Seki M, Shinozaki K, Obokata J (2009) Heterogeneity of Arabidopsis core promoters revealed by high-density TSS analysis. Plant J 60: 350–362

61. Yu G, Wang L-G, He Q-Y (2015) ChIPseeker: an R/Bioconductor package for ChIP peak annotation, comparison and visualization. Bioinformatics 31: 2382–2383

62. Yuan C, Meng X, Li X, Illing N, Ingle RA, Wang J, Chen M (2017) PceRBase: Adatabase of plant competing endogenous RNA. Nucleic Acids Res 45: D1009–D1014

63. Zhang B, Horvath S (2005) AGeneral Framework for Weighted Gene Co-Expression Network Analysis. Stat Appl Genet Mol Biol. doi: 10.2202/1544-6115.1128

64. Zhang Y-C, Liao J-Y, Li Z-Y, Yu Y, Zhang J-P, Li Q-F, Qu L-H, Shu W-S, Chen Y-Q (2014) Genome-wide screening and functional analysis identify a large number of long noncoding RNAs involved in the sexual reproduction of rice. Genome Biol 15: 512

65. Zhang Y, Liu T, Meyer CA, Eeckhoute J, Johnson DS, Bernstein BE, Nusbaum C, Myers RM, Brown M, Li W, et al (2008) Model-based analysis of ChIP-Seq (MACS). Genome Biol 9: R137

66. Zhao T, Tao X, Feng S, Wang L, Hong H, Ma W, Shang G, Guo S, He Y, Zhou B, et al (2018a) LncRNAs in polyploid cotton interspecific hybrids are derived fromtransposon neofunctionalization. Genome Biol 19: 195

67. Zhao X, Li J, Lian B, Gu H, Li Y, Qi Y (2018b) Global identification of Arabidopsis lncRNAs reveals the regulation of MAF4 by a natural antisense RNA. Nat Commun 9: 5056

